# Serotype-dependent bioenergetic and electrophysiological remodelling in iPSC-derived cardiomyocytes from rheumatoid arthritis patients

**DOI:** 10.64898/2026.05.27.727798

**Authors:** Jan Wolnik, Patrycja Adamska, Aleksandra Oleksy, Anna Magdalena Sanetra, Katarzyna Palus-Chramiec, Marian Henryk Lewandowski, Józef Dulak, Monika Biniecka

**Affiliations:** Department of Medical Biotechnology, Faculty of Biochemistry, Biophysics and Biotechnology, Jagiellonian University, Kraków, Poland; Jagiellonian University, Doctoral School of Exact and Natural Sciences, Kraków, Poland; Department of Neurophysiology and Chronobiology, Institute of Zoology and Biomedical Research, Jagiellonian University, Kraków, Poland; Silesian Park of Medical Technology, Kardio-Med Silesia, Zabrze, Poland

**Keywords:** rheumatoid arthritis, induced pluripotent stem cells, cardiomyocytes, bioenergetics, electrophysiology

## Abstract

**Background:** Patients with rheumatoid arthritis (RA) exhibit increased cardiovascular morbidity and mortality that are not fully explained by traditional risk factors, with cardiovascular outcomes differing between seropositive (spRA) and seronegative (snRA) disease. The cellular mechanisms linking chronic inflammation to cardiac dysfunction remain poorly defined, and no patient-specific cardiomyocyte model has resolved cellular phenotypes by RA serotype.

**Methods:** Human induced pluripotent stem cell-derived cardiomyocytes (hiPSC-CMs) were generated from healthy donors and patients with spRA and snRA. Bioenergetic and electrophysiological responses to key RA proinflammatory cytokines (TNF-α, IL-1β, IL-6) and anti-rheumatic drugs (adalimumab, tofacitinib) were assessed using Seahorse assays, patch-clamp electrophysiology, multi-electrode array recordings and RT-qPCR.

**Results:** snRA cardiomyocytes exhibited impaired TNF-α-induced oxidative phosphorylation, accompanied by attenuated expression of *ATP5B*, *LDHA* and *DLD*. In contrast, spRA cardiomyocytes showed baseline electrophysiological alterations, including shortened APD90 and increased action-potential triangulation. TNF-α depolarised the maximum diastolic potential in both RA serotypes. At the multicellular level, cytokine effects were serotype-specific: IL-1β selectively prolonged QT interval in spRA monolayers (p < 0.001), whereas IL-6 prolonged QT in snRA (p < 0.05). Both RA serotypes showed impaired TNF-α-driven induction of *KCNJ3* and *KCNA5*. Adalimumab selectively induced *ATP5B* in spRA but failed to engage either pathway in snRA, while tofacitinib selectively induced *KCNJ3* in healthy but not RA cardiomyocytes.

**Conclusions:** These findings define distinct, serotype-specific pathways of cardiac remodelling in RA that converge on a shared proarrhythmic phenotype, provide a cellular framework for cardiovascular risk in RA and identify candidate mechanisms relevant to therapy-associated cardiovascular safety.

**Graphical Abstract:** 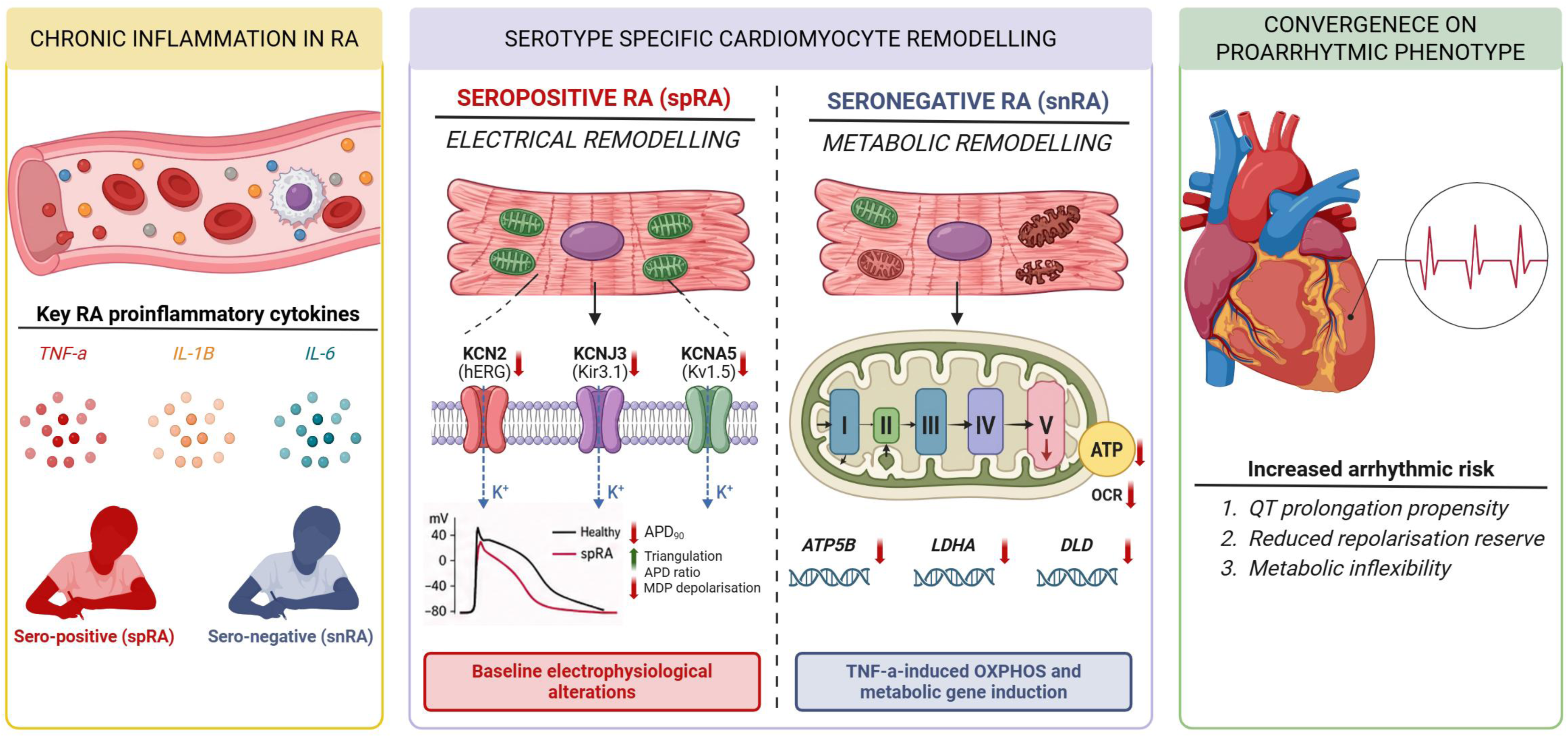

## 1. Introduction

Rheumatoid arthritis (RA) is a chronic, systemic autoimmune disease affecting approximately 0.5–1% of the global population. Although joint destruction remains its most visible manifestation, cardiovascular disease (CVD), including atherosclerosis, heart failure, conduction abnormalities and sudden cardiac death, is now recognized as the leading cause of premature mortality in RA [1,2]. Epidemiological studies indicate that patients with RA exhibit up to a twofold increased risk of myocardial infarction and a significantly higher incidence of arrhythmias and heart failure compared with the general population [3]. Importantly, this excess cardiovascular risk cannot be fully explained by traditional risk factors, pointing towards chronic systemic inflammation as a central driver of cardiac dysfunction [1,4]. Population-based studies have further linked RA with QT interval prolongation and increased arrhythmic risk, suggesting a direct effect of inflammatory mediators on cardiomyocyte function [4,5].

RA is a heterogeneous disease comprising distinct clinical and immunological subtypes, most prominently seropositive (spRA) and seronegative (snRA) RA. Seropositive RA, defined by the presence of rheumatoid factor and/or anti-citrullinated protein antibodies, is generally associated with a higher inflammatory burden, more aggressive disease course, and increased systemic complications compared with seronegative RA [6,7]. In contrast, seronegative RA often presents with a more variable clinical phenotype and may involve distinct underlying mechanisms. These subtype-specific differences are increasingly recognized at the immunological and molecular levels [8].

A hallmark of RA is the sustained overproduction of pro-inflammatory cytokines, particularly tumour necrosis factor alpha (TNF-α), interleukin-1 beta (IL-1β), and interleukin-6 (IL-6), which form a self-amplifying inflammatory network. These mediators exert systemic effects extending beyond the joint microenvironment and have been implicated in endothelial dysfunction, vascular remodelling, and myocardial injury [4,9]. Their circulating levels may persist for years in active disease and differ between clinical subtypes, with spRA typically characterised by higher TNF-α and IL-6 levels than snRA. Increasing evidence suggests that circulating cytokines can directly influence cardiac electrophysiology, with TNF-α, IL-1β and IL-6 linked to QT interval prolongation and arrhythmogenic risk [10,11].

One of the emerging concepts in chronic inflammatory diseases is metabolic reprogramming as a driver of cellular dysfunction. In RA, multiple cell types exhibit a shift towards increased glycolysis accompanied by impaired mitochondrial oxidative phosphorylation [12–14]. Mitochondria are central regulators of cardiomyocyte function, not only through ATP production but also via control of reactive oxygen species (ROS), calcium homeostasis, and cell survival pathways. Pro-inflammatory cytokines have been shown to induce mitochondrial dysfunction, characterised by increased ROS generation, disruption of mitochondrial membrane potential (ΔΨm), and structural damage to mitochondrial components [15,16]. These alterations may have direct electrophysiological consequences, as redox imbalance and mitochondrial depolarisation are known to modulate ion channel activity and action potential dynamics.

In cardiomyocytes, electrophysiological properties are tightly coupled to metabolic state. Ion channel function, including sodium (INa), calcium (ICa,L), and potassium currents (IKr, IKs), depends on cellular energy availability and redox homeostasis [17]. Disruption of these processes can lead to delayed repolarisation, reflected clinically as QT interval prolongation, and increase susceptibility to arrhythmias. Despite growing clinical evidence linking inflammation with electrical abnormalities [5,18], direct mechanistic studies in human cardiomyocytes remain limited due to restricted access to primary cardiac tissue and the limitations of animal models, which do not fully recapitulate human cardiac electrophysiology.

Human induced pluripotent stem cell-derived cardiomyocytes (hiPSC-CMs) provide a scalable, patient-specific platform to overcome these limitations. These cells recapitulate key electrophysiological and metabolic features of human cardiomyocytes while enabling controlled exposure to defined inflammatory and pharmacological stimuli [19,20]. Although widely used to model inherited cardiomyopathies and channelopathies [21,22], their application to inflammation-driven cardiac dysfunction in autoimmune diseases remains underexplored, particularly in the context of RA.

The expanding therapeutic landscape in RA, including TNF-neutralising monoclonal antibodies such as adalimumab (ADA) and Janus kinase (JAK) inhibitors such as tofacitinib (TOFA), has significantly improved disease outcomes but has also raised new questions regarding cardiovascular safety [23,24]. While these agents effectively modulate systemic inflammation, their direct effects on cardiomyocyte bioenergetics and electrophysiology remain unclear, especially in a patient-specific human cellular context.

In this study, we generated hiPSC-derived cardiomyocytes from healthy donors as well as patients with seropositive and seronegative RA and comprehensively characterized their bioenergetic and electrophysiological responses to TNF-α, IL-1β, IL-6, adalimumab, and tofacitinib. By integrating metabolic and electrophysiological analyses, this study aimed to provide mechanistic insight into how chronic inflammation contributes to cardiac abnormalities in RA.

## 2. Methods

Please refer to the Supplementary Methods for a detailed description of the experimental procedures used in this study.

### 2.1. Patient recruitment and ethical approvals

Whole blood was collected from 3 healthy donors at the Silesian Park of Medical Technology, Kardio-Med Silesia, Zabrze, Poland (Bioethical Committee approval no. 3/2019), and from 8 patients with active rheumatoid arthritis at the Clinical Department of Rheumatology and Internal Medicine in Wrocław, Poland (no. KB 111/2020). Synovial biopsies were obtained from RA patients undergoing knee arthroscopy at the Rheumatology Department, St Vincent’s University Hospital, Dublin, Ireland; fibroblast-like synoviocytes (FLS) isolated from these biopsies were kindly provided by Prof. Ursula Fearon (Trinity College Dublin, Ireland). Eight RA patients were enrolled: 4 with seropositive RA (n = 4; 2 PBMC-derived and 2 FLS-derived hiPSC lines) and 4 with seronegative RA (n = 4; 2 PBMC-derived and 2 FLS-derived hiPSC lines). hiPSC lines from healthy donors were generated exclusively from PBMCs (n = 3). All patients fulfilled the 2010 ACR/EULAR classification criteria for RA, had high disease activity (DAS28 > 5.1) and had not received prior biologic therapy. Seropositivity was defined by the presence of rheumatoid factor and/or anti-citrullinated protein antibodies. All procedures were conducted in accordance with the Declaration of Helsinki, and written informed consent was obtained from all participants. Donor demographic and clinical characteristics are summarized in Supplementary Table 1.

### 2.2. Generation, characterization and cardiac differentiation of hiPSC lines

hiPSC lines were generated from PBMCs and FLS by Sendai virus-mediated reprogramming using the CytoTune-iPS 2.0 Sendai Reprogramming Kit (Thermo Fisher Scientific), following established protocols [25]. IPSC lines were validated by karyotyping (G-banding GTG-450) (Kariogen Laboratory, Kraków, Poland), pluripotency-marker expression (immunofluorescence) and trilineage differentiation potential (*in vitro* embryoid body formation and *in vivo* teratoma assay in NOD SCID mice). Animal procedures were approved by the Second Local Ethical Committee for Animal Research in Kraków (approval no. 131/2021) and complied with Directive 2010/63/EU. Cardiac differentiation followed temporal modulation of the Wnt/β-catenin pathway [26]. hiPSCs were sequentially treated with the GSK3β inhibitor CHIR-99021 (6 µM, day 0) and the Wnt antagonist IWR-1 (4–7 µM, line-optimized, day 3) in RPMI-1640 supplemented with 2% B-27 minus insulin. From day 7 onwards, complete RPMI/B-27 medium was used. On day 21, monolayers were enriched to >90% cTnT^+^ purity by magnetic-activated cell sorting (PSC-Derived Cardiomyocyte Isolation Kit, Miltenyi Biotec) [27]. All functional assays were performed between days 30 and 45 post-differentiation.

### 2.3. Stimulation of hiPSC-derived cardiomyocytes

hiPSC-CMs were stimulated for 24 h in RPMI-1640/B-27 with TNF-α (10 ng/mL), IL-1β (1 ng/mL) or IL-6 (20 ng/mL) (all Bio-Techne/R&D Systems), or with tofacitinib (10 µM) or adalimumab (10 mg/mL) (both MedChemExpress). Concentrations were selected based on preliminary viability assays and previously published protocols [28–31]. Paired unstimulated wells from the same differentiation batch served as internal controls for each patient-specific line.

### 2.4. Mitochondrial function and bioenergetics

Mitochondrial function was assessed in MACS-purified hiPSC-CMs (100,000 cells/well) by flow cytometry (LSR Fortessa, BD Biosciences) using MitoTracker Green FM, MitoSOX Red, CellROX Deep Red, tetramethylrhodamine methyl ester (TMRM, ΔΨm) and JC-1 (ΔΨm by aggregate/monomer ratio). Tert-butyl hydroperoxide and FCCP served as positive controls. Subcellular probe localisation was confirmed by live-cell holotomography (HT-X1, Tomocube). Cellular bioenergetics was profiled using the Seahorse XF Mito Stress Test (Agilent). Purified hiPSC-CMs were plated at 15,000 cells/well on Geltrex-coated XF96 plates and cultured for 72 h before stimulation. Oxygen consumption rate (OCR) was recorded after sequential injections of oligomycin (1 µg/mL), FCCP (1 µM) and rotenone/antimycin A (1 µM each). Basal respiration, ATP-linked respiration, maximal respiration and spare respiratory capacity were calculated using Wave software (Agilent) and expressed as % of paired unstimulated controls.

### 2.5. Gene expression analysis

Total RNA was extracted using the RNeasy Mini Kit (Qiagen). cDNA was generated from 0.5 µg RNA using the High-Capacity RNA-to-cDNA Kit (Thermo Fisher). Real-time PCR was performed using AceQ Universal SYBR qPCR Master Mix (Vazyme) on a StepOnePlus thermocycler. Expression of metabolic (*ATP5B*, *LDHA*, *DLD*) and ion-channel (*KCNH2*, *KCNA5*, *KCNJ3*, *SCN5A*, *CACNA2*) transcripts were normalised to the cardiac-specific reference gene *TNNT2*.

### 2.6. Single-cell patch-clamp electrophysiology

MACS-purified hiPSC-CMs were replated at a density of 7,500 cells per 10-mm Geltrex-coated round glass coverslip, which was maintained in a 24-well plate during coating and cell seeding. Three days after replating, cells were stimulated for 24 h with cytokines as above. Recordings were performed in whole-cell current-clamp configuration at 37 ± 0.5 °C using an SC 05LC amplifier (NPI Electronic), low-pass filtered at 2 kHz, digitised at 20 kHz and acquired with Signal software (Cambridge Electronic Design). Borosilicate glass electrodes (tip resistance 7–9 MΩ) were filled with intracellular solution containing (mM): 125 K-gluconate, 20 KCl, 5 NaCl, 10 HEPES (pH 7.2 with KOH). Cells were superfused with extracellular solution containing (mM): 140 NaCl, 5.4 KCl, 1.8 CaCl_2_, 1.0 MgCl_2_, 5.5 glucose, 5 HEPES (pH 7.4 with NaOH). Resting membrane potential was manually adjusted to −60 mV by continuous negative current injection. Single action potentials were elicited at 1 Hz using brief rectangular depolarising current pulses (2 ms, 150–210 pA, adjusted per cell). Ten consecutive action potentials per cell were averaged for parameter extraction. Cells were excluded if series resistance exceeded 20 MΩ or changed by more than 20% during recording, AP amplitude was below 70 mV, or pacemaker-like phase 4 depolarisation was observed. Six AP parameters were extracted: maximum upstroke velocity (dV/dt_max_), AP peak amplitude, AP through, APD20, APD90 and APD ratio (30–40 / 70–80), (see Supplementary Table 4 for definitions and physiological interpretation of action potential parameters). Recordings were obtained from N = 3 donors, with n > 20 cells per group across independent differentiation batches. Analyses were performed blinded to donor group identity.

### 2.7. Multi-electrode array recordings

Spontaneous field potentials were recorded from hiPSC-CM monolayers seeded on Geltrex-coated 24-well MEA plates (Multi Channel Systems) using the Multiwell-MEA system at 37 °C. After 4 days of culture allowing monolayer formation, baseline field potential activity was recorded for 5 min (sampling 20 kHz; 1 Hz high-pass and 3.5 kHz low-pass filters). Cytokines and drugs were applied as 5× concentrated stimulants and recordings were repeated after 24 h with identical settings. QT-equivalent interval, beat period and beat frequency were extracted using Multiwell-MEA-Analyser software with manual verification. Each well was normalised to its own baseline and compared with paired unstimulated controls; minimum three wells per condition with at least 3 electrodes capturing optimal filed potential signal.

### 2.8. Statistical analysis

Data are expressed as mean ± standard deviation (SD) from at least three independent biological replicates, unless otherwise stated. Normality was assessed using the Shapiro–Wilk test. Group comparisons used the Kruskal–Wallis test with Dunn’s post-hoc correction (non-parametric), or one-way ANOVA with Bonferroni correction for normally distributed data. Two-way ANOVA with Dunnett’s post-hoc test was applied when both donor group and stimulation condition were assessed simultaneously. Analyses were performed in GraphPad Prism 10. A two-tailed *p* < 0.05 was considered statistically significant. Significance levels: \**p* < 0.05; \*\**p* < 0.01; \*\*\**p* < 0.001; \*\*\*\**p* < 0.0001.

## 3. Results

### 3.1. Generation and characterization of patient-derived hiPSCs and hiPSC-CMs

An overview of the experimental workflow is presented in Fig. 1. PBMCs and FLS obtained from HC, spRA and snRA donors were successfully reprogrammed into hiPSCs using Sendai virus vectors. All generated lines exhibited typical pluripotent stem cell morphology with compact colonies and well-defined borders (Fig. 2). G-banded karyotype analysis (GTG-450, minimum 20 metaphases per line) confirmed a normal karyotype with no numerical or structural aberrations (Supplementary Fig. 1A). Immunofluorescence analysis confirmed expression of key pluripotency markers, including SSEA-4, OCT-4, NANOG, TRA-1-60 and TRA-81 (Fig. 2 and Supplementary Fig. 1B). Embryoid body formation assays demonstrated capacity for spontaneous differentiation, evidenced by lineage-specific markers including NFH (ectoderm). Teratoma formation assays confirmed *in vivo* pluripotency with histological analysis revealing tissues representative of all three germ layers (Fig. 2). Next, hiPSCs were differentiated into ventricular-like cardiomyocytes via temporal Wnt-pathway modulation and differentiation efficiency was evaluated by intracellular flow cytometry for cardiac troponin T (cTnT) on day 21. Across donor lines, 30–80% of live single cells were cTnT⁺ prior to MACS sorting (representative gating in Supplementary Fig. 1C). Cardiomyocytes were subsequently enriched to >80% cTnT⁺ purity by MACS sorting before downstream functional assays. Resulting hiPSC-CMs displayed characteristic morphology, forming spontaneously contracting clusters and organized monolayers. Immunofluorescence confirmed positive staining for the cardiomyocyte markers MYL2, MYH7 and ACTN2 with well-defined sarcomeric organization, together with the gap-junction protein Cx43 and the cardiac transcription factors NKX2.5 and HOPX. (Fig. 2).

**Figure 1.**
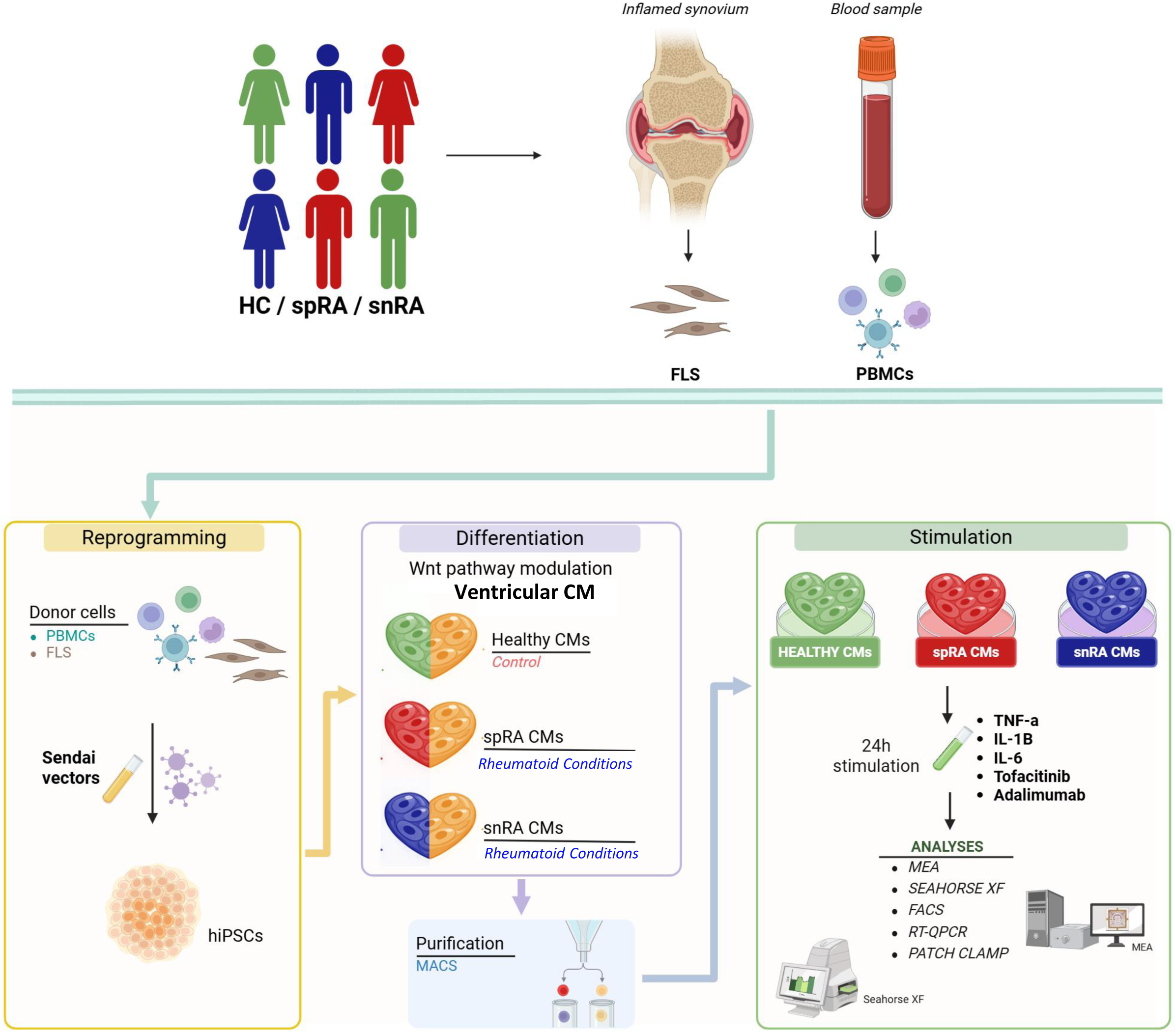
Experimental overview and study design. Schematic representation of the experimental workflow. Peripheral blood mononuclear cells (PBMCs) and fibroblast-like synoviocytes (FLS) obtained from healthy controls (HC), seropositive RA (spRA), and seronegative RA (snRA) donors were reprogrammed into hiPSCs using Sendai virus vectors. hiPSCs were subsequently differentiated into ventricular-like cardiomyocytes via temporal modulation of the Wnt signalling pathway and purified by magnetic-activated cell sorting (MACS). Purified hiPSC-CMs were exposed for 24 h to TNF-α, IL-1β, IL-6, tofacitinib (TOFA), or adalimumab (ADA), followed by functional and molecular analyses using Seahorse XF, flow cytometry, RT-qPCR, whole-cell patch-clamp electrophysiology, and multi-electrode array (MEA) platforms.

**Fig. 2.**
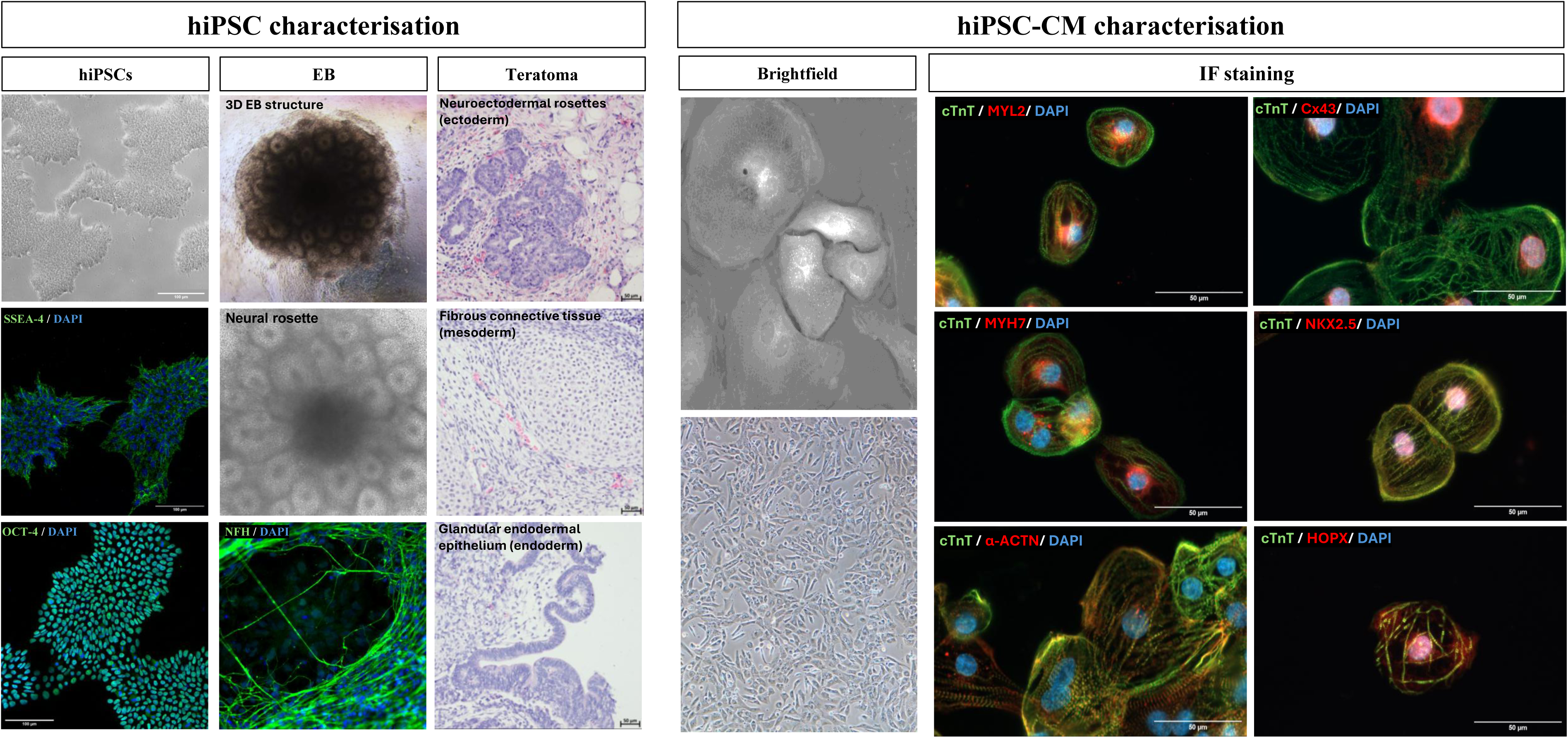
Characterization of patient-derived hiPSCs and hiPSC-CMs. *(Left) hiPSC characterization*. Brightfield images of typical undifferentiated hiPSC colonies and immunofluorescence staining for pluripotency markers SSEA-4 (green) and OCT4 (green), with nuclei counterstained with DAPI (blue). Embryoid body (EB) formation demonstrates three-dimensional structures and neural rosette formation, with NFH (green) confirming neuroectodermal differentiation. Teratoma formation is shown by representative haematoxylin and eosin-stained sections containing tissues from all three germ layers. ***(Right) hiPSC-CM characterization***. Brightfield images show typical morphology of spontaneously beating cardiomyocyte clusters and confluent monolayers. Immunofluorescence staining for cTnT (green), MYL2, Cx43, HOPX, NKX2-5, ACTN2 and MYH7 (all red), with well-defined sarcomeric organisation. Nuclei were counterstained with DAPI (blue).

### 3.2. Mitochondrial homeostasis is preserved across donor groups

To assess the effects of pro-inflammatory cytokines and anti-rheumatic drugs on mitochondrial homeostasis, three parameters were quantified in parallel cultures: cellular ROS, mitochondrial mass and mitochondrial membrane potential (ΔΨm), assessed using complementary probes (TMRM and JC-1) (Fig. 3). Following 24 h stimulation, fluorescence readouts were normalized to paired unstimulated controls. Intracellular ROS (CellROX Deep Red) did not differ across treatments or donor groups, while the positive control TBHP induced a clear increase (Fig. 3A). Mitochondrial mass (MitoTracker Green) remained unchanged (Fig. 3B). ΔΨm measured by TMRM was stable across all conditions, whereas FCCP induced a marked reduction (Fig. 3C), a finding confirmed by JC-1, where only FCCP-treated controls showed a significant shift in red/green fluorescence ratio (Fig. 3D). Live-cell holotomography combined with multichannel fluorescence (Fig. 3E) demonstrated co-localisation of MitoTracker and MitoSOX signals throughout the mitochondrial network, with substantial overlap of CellROX signal in mitochondrial regions. Importantly, these holotomographic and fluorescence microscopy observations complement the flow cytometry data by providing spatial validation of the detected fluorescence signals and confirming that the ROS signals quantified in bulk flow cytometry analyses originate primarily from the mitochondrial compartment. Collectively, these results indicate that, under the tested conditions, pro-inflammatory cytokines and anti-rheumatic treatments do not alter mitochondrial ROS levels, membrane potential or mitochondrial content, irrespective of RA serotype.

**Fig. 3.**
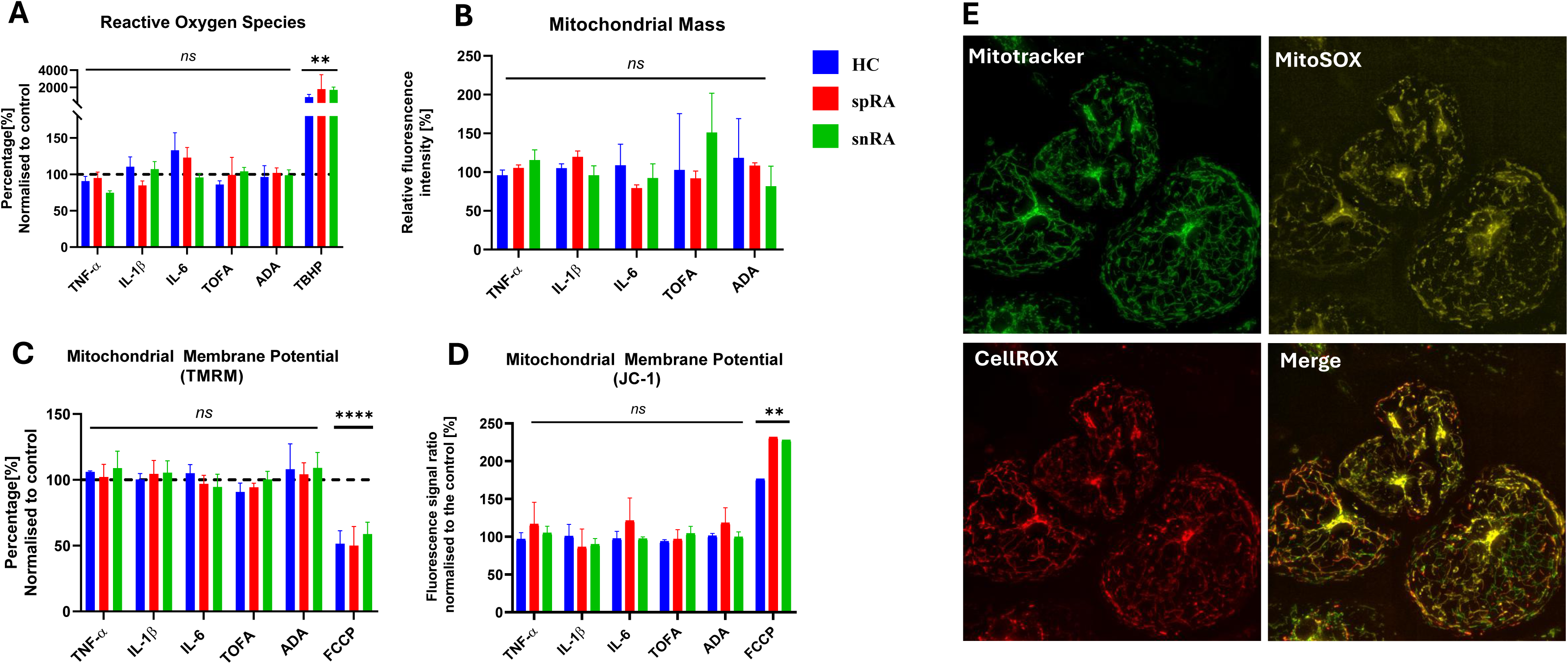
Mitochondrial properties and ROS distribution in hiPSC-CMs. Quantitative assessment of mitochondrial function and oxidative status in hiPSC-CMs derived from healthy controls (HC, blue), seropositive RA (spRA, red), and seronegative RA (snRA, green) following 24 h stimulation. All values are normalized to paired unstimulated controls (dashed line at 100%). **(A)** Intracellular ROS measured by CellROX Deep Red using flow cytometry; tert-butyl hydroperoxide (TBHP) served as a positive control. **(B)** Mitochondrial mass assessed by MitoTracker Green fluorescence. **(C)** Mitochondrial membrane potential (ΔΨm) measured by TMRM; FCCP served as a positive control for depolarisation. **(D)** ΔΨm assessed by JC-1 red/green fluorescence ratio; FCCP served as a positive control. **(E)** Live cell holotomographic imaging combined with multichannel fluorescence showing mitochondrial network and ROS distribution. Mitochondria were labelled MitoTracker Green, mitochondrial superoxide with MitoSOX Red and intracellular ROS with CellROX Deep Red; merged images illustrate spatial overlap of signals. Images were acquired at 40× magnification using an HT-X1 holotomography microscope (Tomocube). Data are presented as mean ± SD from at least three independent experiments. Statistical analysis was performed using the Kruskal–Wallis test with Dunn’s post-hoc correction. **p < 0.01, **p < 0.0001; ns, not significant.

### 3.3. Divergent metabolic and transcriptional responses define serotype-specific vulnerability in rheumatoid arthritis cardiomyocytes

Seahorse mitochondrial stress tests revealed a serotype-specific bioenergetic response (Fig. 4A–D). Neither IL-1β nor IL-6 induced consistent changes in mitochondrial respiration in any donor group, whereas TNF-α elicited a strongly divergent response. In HC and spRA cardiomyocytes, TNF-α triggered an adaptive increase in mitochondrial respiration, while in snRA cardiomyocytes this response was absent, with OCR values remaining below baseline across all measured parameters. This divergence, confined to TNF-α and not observed with IL-1β or IL-6, identifies TNF-α as the central driver of bioenergetic remodelling under these conditions. Tofacitinib did not affect mitochondrial respiration in any donor group, whereas adalimumab induced an increase in mitochondrial function selectively in HC cardiomyocytes, with no comparable response in either RA serotype, suggesting a loss of baseline sensitivity to TNF signalling in RA. Consistent with these bioenergetic findings, transcriptional analysis (Fig. 4E–G) revealed reduced adaptability of metabolic genes in RA cardiomyocytes. TNF-α induced coordinated upregulation of *ATP5B*, *LDHA* and *DLD* in HC cells, whereas this response was markedly attenuated in both RA serotypes. IL-1β elicited a weaker, HC-dominant response for *ATP5B* and *LDHA*, while IL-6 had no significant effect on any of the three transcripts. Adalimumab selectively induced *ATP5B* in spRA cardiomyocytes, whereas tofacitinib induced *DLD* in HC without affecting the other transcripts. Collectively, these findings establish TNF-α as the central cytokine driving bioenergetic and transcriptional remodelling in hiPSC-CMs and reveal a serotype-specific vulnerability profile, where seronegative RA cardiomyocytes lack the adaptive oxidative-phosphorylation response observed in healthy and seropositive cells.

**Fig. 4.**
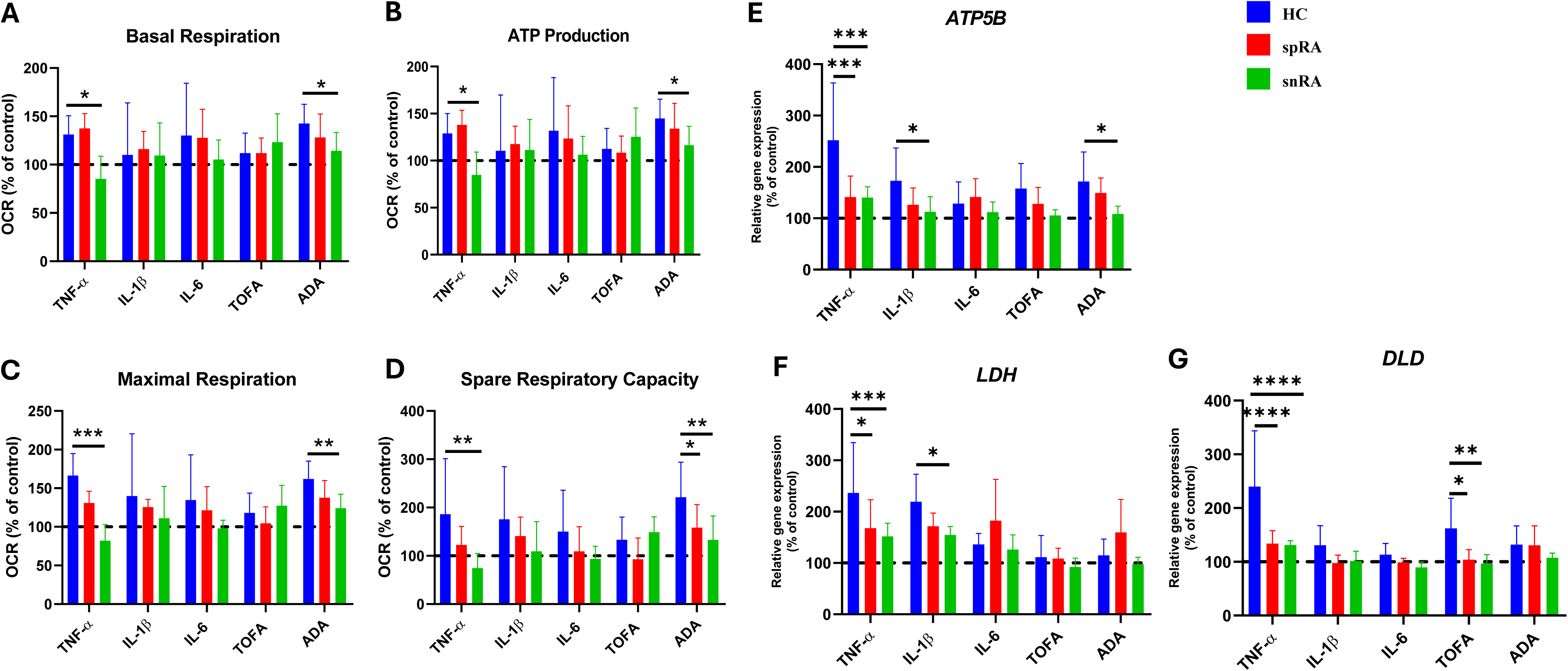
Bioenergetic profiling and metabolic gene expression in hiPSC-derived cardiomyocytes. **(A–D)** Mitochondrial respiration parameters assessed using the Seahorse XF Mito Stress Test in hiPSC-CMs derived from healthy controls (HC, blue), seropositive RA (spRA, red) and seronegative RA (snRA, green): **(A)** basal respiration, **(B)** ATP-linked respiration, **(C)** maximal respiration, and **(D)** spare respiratory capacity. Oxygen consumption rate (OCR) is expressed as a percentage of the paired unstimulated control for each donor line. **(E–G)** Relative expression of metabolism-related genes measured by RT-qPCR: **(E)** *ATP5B*, **(F)** *LDHA* and **(G)** *DLD*. Gene expression levels were normalized to *TNNT2* and *EEF2* and are presented relative to paired unstimulated controls. Data are shown as mean ± SD from 2–3 independent differentiations per donor (n = 3 HC, n = 3 spRA, n = 3 snRA donors). Statistical analysis was performed using the Kruskal–Wallis test with Dunn’s post-hoc correction. *p < 0.05; **p < 0.01; ***p < 0.001; ***p < 0.0001.

### 3.4. Patch-clamp electrophysiology reveals serotype-specific remodelling of action potential properties

To define the cellular basis of cytokine-induced electrical remodelling at the single-cell level, we performed whole-cell patch-clamp recordings under basal conditions and following 24 h exposure to TNF-α, IL-1β or IL-6, quantifying six action potential parameters per averaged trace (dV/dtmax, peak amplitude, APD20, APD90, APD ratio and AP through) (Fig. 5A and Supplementary Table 4). APD90 was consistently shortened in both RA serotypes across all conditions, indicating constitutive remodelling of late repolarisation (Fig. 5B top panel). In contrast, APD20 exhibited a stimulus-dependent, serotype-selective pattern, with shortening observed in snRA under basal and IL-1β conditions and in spRA following TNF-α exposure, while IL-6 had no measurable effect (Fig. 5B middle panel). Further analysis of action potential morphology revealed a marked increase in APD ratio (30–40 / 70–80) in spRA cardiomyocytes across all conditions, consistent with a constitutive shift towards triangulated action potentials, whereas snRA showed a similar but weaker pattern only under stimulation. TNF-α further induced selective depolarisation of the maximum diastolic potential in both RA serotypes, an effect not observed with IL-1β or IL-6 (Fig. 5B bottom panel). Given that AP through is principally governed by the inward rectifier potassium currents IK1 (encoded by *KCNJ2/KCNJ12*) and IK,ACh (encoded by *KCNJ3*), this TNF-α-selective depolarisation provides a direct functional correlate of the loss of *KCNJ3* induction observed at the transcriptional level (Supplementary Fig. 2 bottom panel). In contrast, neither upstroke velocity nor action potential amplitude showed consistent differences between groups, localizing the observed phenotype to potassium-dependent repolarisation rather than depolarisation mechanisms (Supplementary Fig. 2 top and middle panel respectively). Together, these single-cell findings localize the RA-associated electrical phenotype to potassium-dependent repolarisation, with constitutive APD90 shortening in both serotypes, spRA-specific action potential triangulation and TNF-α-selective AP through depolarisation as three independent signatures pointing to a serotype-shared loss of *KCNJ3* induction.

**Fig. 5.**
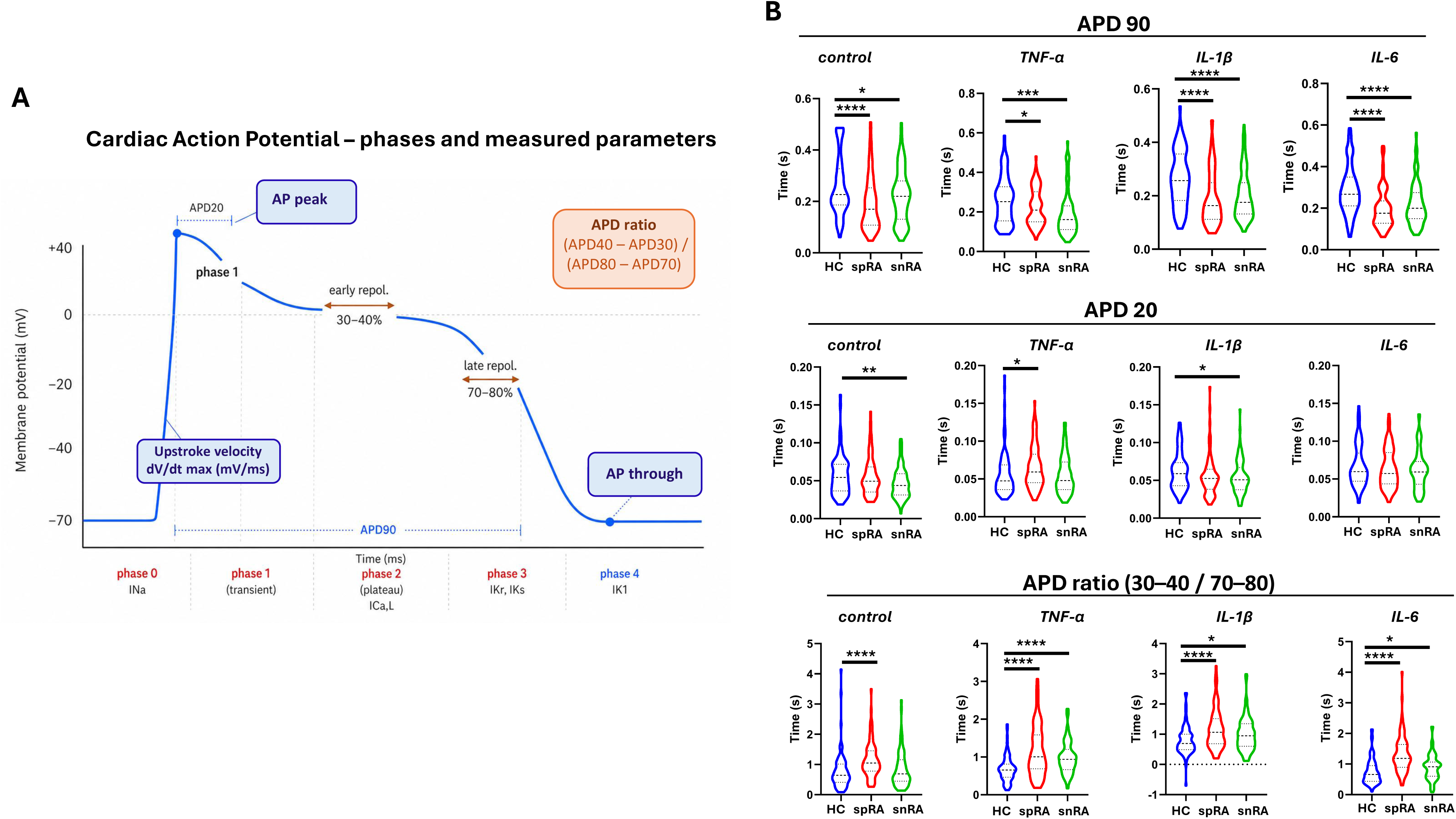
Single-cell patch-clamp electrophysiology reveals serotype-specific remodelling of action potential properties in RA cardiomyocytes. **(A)** Schematic representation of a cardiac action potential (AP) with the five canonical phases (0–4) indicated along the time axis. Six parameters were extracted from averaged traces of 10 consecutive APs: maximum upstroke velocity (dV/dt), AP peak amplitude, action potential duration at 20% and 90% repolarisation (APD20, APD90), APD ratio (30–40 / 70–80) and maximum diastolic potential (MDP). **(B)** Action potential parameters in hiPSC-CMs derived from healthy controls (HC, blue), seropositive RA (spRA, red), and seronegative RA (snRA, green). Top: APD90. Middle: APD20. Bottom: APD ratio (30–40 / 70–80). Plots display the distribution of individual cell measurements, with median (solid line) and interquartile range (dashed lines). Data are shown from n = 3 donors per group. Statistical analysis was performed using the Kruskal–Wallis test with Dunn’s post-hoc correction. *p < 0.05; **p < 0.01; ***p < 0.001; ***p < 0.0001.

### 3.5. MEA recordings reveal a distinct serotype-specific cytokine selectivity at the multicellular level

To extend these single-cell observations to the multicellular level, where cell-to-cell electrical coupling shapes the integrated electrical response, we recorded field potentials from spontaneously beating hiPSC-CM monolayers using multi-electrode array (MEA) technology (Fig. 6A-C). In HC cardiomyocytes, none of the tested cytokines or anti-rheumatic agents altered the QT interval (Fig. 6A). In contrast, spRA cardiomyocytes exhibited a pronounced QT prolongation in response to IL-1β, whereas TNF-α, IL-6, tofacitinib and adalimumab had no measurable effect. In snRA cardiomyocytes, IL-6 induced a modest QT prolongation, while all other conditions remained unchanged. Notably, TNF-α did not affect QT interval in any group despite producing marked changes at the single-cell and transcriptional levels, indicating a dissociation between cellular and multicellular responses. To investigate the molecular basis of this serotype-specific phenotype, we quantified expression of key repolarising potassium channel genes (*KCNH2*, *KCNA5* and *KCNJ3*) (Fig. 6D-F). *KCNH2* showed divergent regulation, with increased expression in HC in response to adalimumab, no change in spRA, and reduced expression in snRA following IL-6 exposure, consistent with the QT prolongation observed in this serotype. In contrast, *KCNA5* and *KCNJ3* displayed a strong HC-dominant pattern, with robust TNF-α-driven induction in HC cardiomyocytes that was absent in both RA serotypes, indicating a shared impairment of cytokine-dependent potassium channel regulation. This loss extended to weaker stimuli, with HC cells maintaining inducibility of *KCNJ3* across conditions, whereas RA cardiomyocytes remained unresponsive. Additionally, spRA cardiomyocytes showed reduced *KCNA5* expression under IL-6, suggesting a serotype-specific layer of repolarisation vulnerability. Finally, tofacitinib selectively induced *KCNJ3* in HC but not in RA cardiomyocytes, further highlighting a loss of adaptive repolarisation reserve in the RA cellular context. Expression of additional ion channel genes, including *CACNA2* and *SCN5A*, remained unchanged across all conditions and donor groups (data not shown). Together, these results show three key features of cardiomyocyte dysfunction in RA at the multicellular level. First, different RA subtypes respond to different cytokines in terms of QT prolongation: IL-1β affects spRA, while IL-6 affects snRA. Second, both RA subtypes lose the ability to activate *KCNA5* and *KCNJ3* in response to cytokines, regardless of autoantibody status. Third, snRA shows a specific reduction of *KCNH2* expression in response to IL-6.

**Fig. 6.**
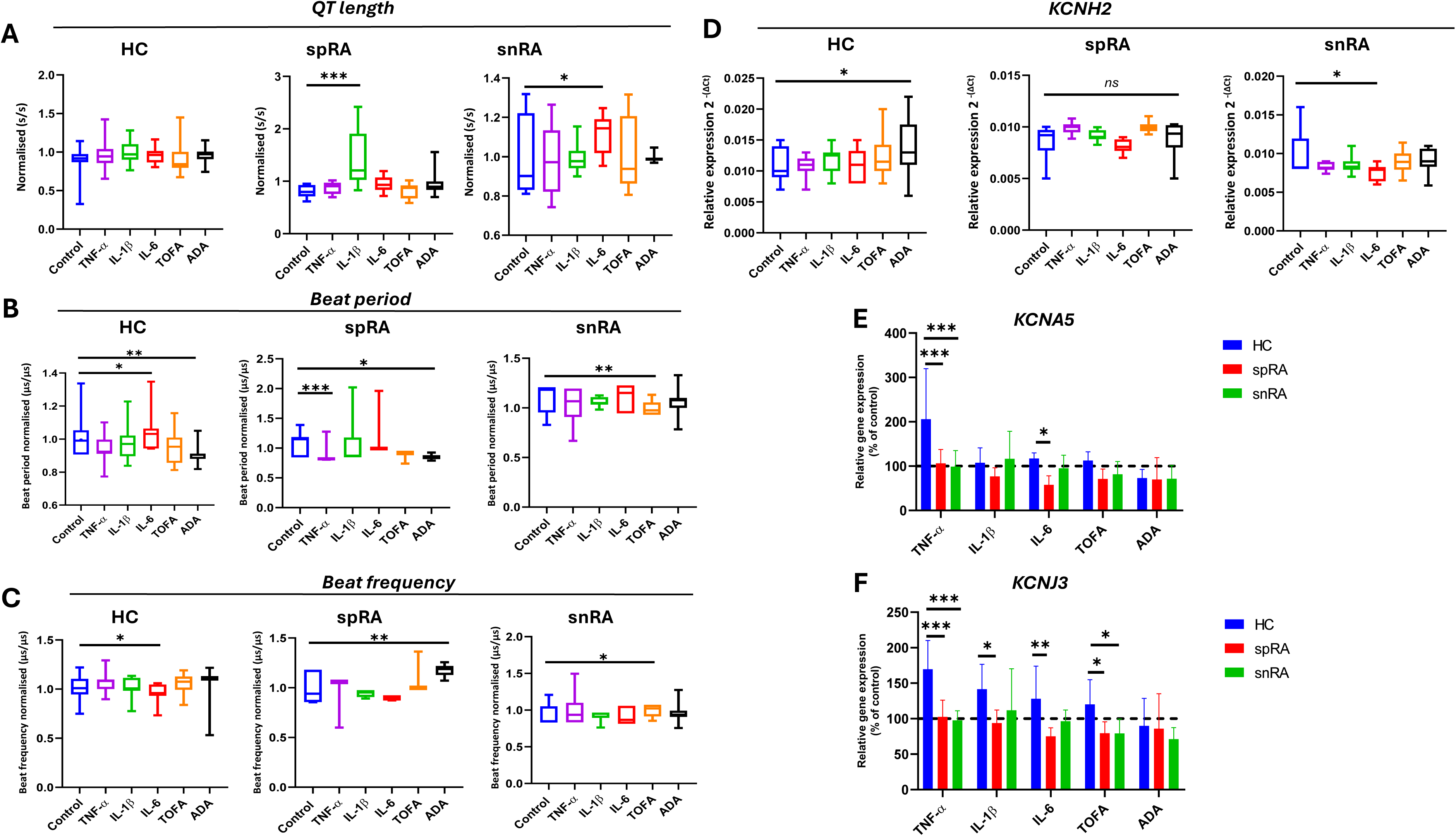
MEA recordings and ion channel gene expression reveal serotype-specific responses in RA cardiomyocytes. MEA (**A–C**): QT-equivalent interval is prolonged by IL-1β in spRA and by IL-6 in snRA, with no changes in HC. Beat period and frequency show condition- and serotype-dependent effects: IL-6 increases beat period and reduces frequency in HC, while ADA shortens beat period in the same group; TNF-α and ADA shorten beat period in spRA, with ADA increasing frequency; TOFA reduces both beat period and frequency in snRA. Gene expression (**D–F**): *KCNH2* is increased by ADA in HC and reduced by IL-6 in snRA, with no change in spRA. *KCNA5* is induced by TNF-α in HC but not in RA and is reduced by IL-6 in spRA. *KCNJ3* is upregulated by inflammatory stimuli in HC but remains unresponsive in RA. Data are mean ± SD or box plots (n ≥ 3). Kruskal–Wallis with Dunn’s post hoc test (*p < 0.05; **p < 0.01; ***p < 0.001; ns).

### 3.6. Serotype-dependent modulation of contractile dynamics

Serotype-dependent modulation of contractile dynamics was assessed by analysing beat period (inter-beat interval) (Fig. 6B) and beat frequency (Fig. 6C). Beat period and beat frequency were normalized to each line’s unstimulated baseline, yielding control values near unity across all donor groups. Baseline beat period was comparable between groups, indicating preserved spontaneous rhythmic activity under unstimulated conditions. Treatment effects were expressed as deviations from this paired baseline within each line. In HC cardiomyocytes, IL-6 induced a moderate prolongation of beat period accompanied by a reduction in beat frequency and adalimumab was associated with a shortening of beat period. In spRA cardiomyocytes, the most pronounced beat-period shortening occurred under TNF-α, with an additional, smaller reduction under adalimumab, while IL-1β and IL-6 showed wide variability without consistent directional change; adalimumab was associated with an increase in beat frequency. In snRA cardiomyocytes, beat period was largely preserved except for a modest but significant shortening under TOFA, which was also reflected in a small reduction of beat frequency. Together, these data reveal that spRA and snRA cardiomyocytes diverge in their pharmacological sensitivity at the contractile level, with TNF-α/adalimumab as principal modulators in spRA and tofacitinib as the principal modulator in snRA paralleling the cytokine selectivity observed for QT prolongation and providing additional cellular evidence for serotype-dependent responses to anti-rheumatic therapy.

## 4. Discussion

This study establishes the first patient-specific hiPSC-derived cardiomyocyte model of rheumatoid arthritis stratified by autoantibody status and provides new insight into the cellular basis of cardiovascular vulnerability in RA. Although previous work has generated cardiomyocytes from RA patients [32,33] or examined TNF-α effects in hiPSC-CMs from healthy donors [28], none has combined serotype stratification with integrated analyses of energy metabolism and electrophysiology in the context of key RA-related cytokines and anti-rheumatic agents.

Our data reveals a consistent pattern of dysfunction emerging across multiple levels. RA hiPSC-CMs reproduce the clinically observed QT prolongation under inflammatory conditions, with clear serotype specificity at the multicellular level - IL-1β in spRA and IL-6 in snRA. In contrast, TNF-α produces marked effects at the single-cell and transcriptional levels without altering QT interval in multicellular recordings, indicating a dissociation between cellular and tissue-level responses. Within this framework, the two serotypes show distinct vulnerability profiles. spRA is dominated by electrical dysfunction, whereas snRA combines metabolic and electrical abnormalities. Despite these differences, both share a common defect - cytokines fail to upregulate the cardioprotective potassium channels *KCNJ3* and *KCNA5*, regardless of autoantibody status. In addition, tofacitinib fails to activate the cytoprotective *KCNJ3* pathway in RA cardiomyocytes, a finding that is potentially relevant to increased cardiovascular risk reported in clinical studies. [24,34]. Although the underlying trajectories differ, both RA serotypes converge on the same endpoint - a depleted repolarisation reserve, accompanied by electrophysiological features such as QT prolongation, AP triangulation, and AP through depolarisation, all of which are established predictors of arrhythmic susceptibility [35,36]. This shared outcome identifies the cytokine–K⁺ channel induction axis as the central mechanism, layered with serotype-specific vulnerabilities — IL-6/*KCNH2* in snRA and IL-1β/repolarisation reserve loss in spRA.

### 4.1. Clinical outcome and serotype-specific cytokine selectivity

The cytokine-specific effects observed in our model closely align with clinical data. In our system, IL-1β prolongs QT in spRA and IL-6 in snRA, consistent with established associations of these cytokines with QTc prolongation in RA cohorts [5,11] and with the higher QTc reported in ACPA-positive patients [37]. Systemic inflammation has been identified as a key driver of arrhythmic risk in RA, linking elevated cytokines with repolarisation abnormalities and increased incidence of atrial fibrillation, ventricular arrhythmias and sudden cardiac death [4,38]. TNF-α, IL-1β and IL-6 have all been independently associated with QT prolongation [5] and population studies confirm an increased prevalence of QTc abnormalities in RA [18]. CV risk is particularly elevated in seropositive patients. ACPA-positive individuals exhibit longer QT intervals than seronegative patients independent of confounders and show increased cardiovascular risk in large cohorts [39]. Therapeutically, IL-6 blockade (e.g. tocilizumab) reduces QTc duration, and circulating IL-6 levels are a strong predictor of QT prolongation [11,40]. Mechanistically, IL-6 directly inhibits hERG channels, alters intercellular coupling and contributes to QT prolongation in inflammatory states [41]. The selective reduction of *KCNH2* expression in snRA cardiomyocytes observed here provides a cellular correlate for these findings. In contrast, spRA sensitivity to IL-1β is consistent with its known effects on potassium currents and arrhythmogenesis [42]. Together, these observations support autoantibody status as a relevant marker for cardiovascular risk stratification and treatment selection in RA.

### 4.2. Mechanistic insights into the cytokine–K⁺ channel induction defect

A central observation across all levels of analysis is the failure of RA cardiomyocytes to activate potassium channels in response to cytokines. This defect is present in both serotypes and provides a common basis for their proarrhythmic phenotype. At the single-cell level, this is reflected by TNF-α–induced depolarisation of the maximum diastolic potential—from ∼−72 mV in controls to ∼−67 mV in RA cells. This parameter is primarily determined by inward rectifier potassium currents (IK1 and IK,ACh), encoded by *KCNJ2/KCNJ12* and *KCNJ3* [43,17]. The inability of TNF-α to induce *KCNJ3* in RA provides a direct explanation for this depolarisation. Even a ∼5 mV shift increases proximity to the activation threshold, promoting spontaneous activity and reducing electrical stability. Similar mechanisms have been described for IL-1β, which suppresses inward rectifier currents and promotes arrhythmogenic activity [42]. The earliest experimental evidence for direct cytokine modulation of cardiac repolarisation came from Li and Rozanski, who demonstrated that recombinant IL-1 alters electrical properties of guinea pig ventricular cells, establishing a foundational precedent for the cytokine–ion channel axis subsequently extended to TNF-α and IL-6 [44]. Importantly, expression of sodium and calcium channel genes (*SCN5A*, *CACNA2*) was preserved across all conditions and donor groups in our system, localising the cytokine-driven transcriptional defect specifically to the repolarising potassium channel axis rather than to depolarisation or plateau phase ionic mechanisms.

In healthy cardiomyocytes, TNF-α induces coordinated upregulation of *KCNJ3* and *KCNA5*, consistent with an early compensatory response. This adaptation is absent in both RA serotypes despite preserved baseline expression. The persistence of this defect through Sendai-mediated reprogramming and cardiac differentiation suggests that RA-associated transcriptional or epigenetic signatures acquired during chronic inflammatory exposure are retained in the cellular memory of the resulting cardiomyocytes, providing a stable cell-intrinsic substrate for the observed phenotype. Evidence from our 3D cardiac microtissue platform supports this concept [45]. RA-derived microtissues exhibited a significantly greater density of CD31⁺ capillary-like networks than healthy controls, even in the absence of exogenous inflammatory stimulation, reproducing the enhanced pro-angiogenic phenotype that is a hallmark of RA in vivo and confirming that disease-associated, cell-intrinsic properties are preserved across cardiac lineages following reprogramming.

Possible mechanisms include epigenetic changes induced by chronic inflammation, signalling exhaustion or altered receptor pathways such as *TNFR1* desensitisation. snRA shows an additional IL-6–dependent vulnerability. IL-6 inhibits *hERG* currents and affects cardiac electrophysiology, while also influencing other channels and mitochondrial pathways [46–48]. The selective downregulation of *KCNH2* in snRA provides a mechanistic basis for IL-6–driven QT prolongation, indicating that IL-6 becomes the dominant regulator of repolarisation in this context. The timing of cytokine exposure also explains the metabolic findings. In our 24-hour model, TNF-α increases mitochondrial respiration in control and spRA cells, likely reflecting an early adaptive response. Longer exposure is known to lead to mitochondrial dysfunction [49,16,28]. Notably, this adaptive phase is absent in snRA, indicating impaired metabolic flexibility.

### 4.3. Multi-scale dissociation of TNF-α effects

A key observation is the divergence between TNF-α effects at the single-cell and multicellular levels. TNF-α produces pronounced electrical and metabolic changes in individual cells but does not alter QT interval in MEA recordings. This discrepancy likely reflects gap junction-mediated electrical coupling within cardiomyocyte networks, which attenuates local changes through current-sink effects. Direct mechanistic support for this interpretation comes from George and colleagues, who demonstrated that TNF-α modulates cardiac conduction by altering connexin 43 (Cx43) expression and gap junction-mediated electrical coupling between myocytes [50], a mechanism that could blunt the propagation of single-cell-level changes to the multicellular field potential. The importance of intercellular crosstalk in shaping iPSC-CM functional readouts has been further established by Giacomelli and colleagues, who showed that Cx43-mediated communication between cardiomyocytes and supporting non-cardiomyocyte cell types is critical for the maturation and integrated electrical behaviour of multicellular cardiac constructs [51]. As a result, substantial single-cell alterations may not affect the overall signal. In contrast, IL-1β and IL-6 likely act through additional mechanisms, including calcium handling and intercellular connectivity, which influence network behaviour more directly.

Previous studies in cardiomyocyte systems have demonstrated stronger TNF-α effects with longer exposure times [16,28], suggesting that our 24-h paradigm captures an early adaptive phase of response before network-level dysfunction emerges. This has important implications for safety assessment. Standard MEA-based assays may underestimate cellular-level dysfunction, particularly in inflammatory contexts. Combining multicellular recordings with single-cell approaches such as patch clamp may provide a more accurate assessment of proarrhythmic risk in disease models.

### 4.4. Metabolic phenotype of seronegative RA

snRA cardiomyocytes show a clear metabolic deficit, characterised by failure to increase oxidative phosphorylation in response to TNF-α. Specifically, all respiratory parameters remain at or below baseline, in contrast to the adaptive upregulation observed in HC and spRA, indicating a loss of metabolic plasticity rather than a baseline mitochondrial defect. The selective loss of spare respiratory capacity in snRA - the parameter most directly reflecting the cell’s reserve to meet sudden energetic demand - indicates that these cardiomyocytes lack the bioenergetic flexibility required to respond to subsequent physiological or pathological stressors, a feature with direct implications for cardiac vulnerability under inflammatory stress. This cellular finding aligns with systemic metabolic alterations recently documented at the patient level by Wu and colleagues, who identified distinct circulating metabolomic profiles distinguishing RA patients from healthy controls and demonstrated that metabolic dysregulation in RA extends well beyond the synovial compartment [52]. This mirrors metabolic alterations described in RA immune and stromal cells, including disrupted energy production in T-cells, altered macrophage metabolism and broader mTOR-driven reprogramming, with mitochondrial dysfunction increasingly recognised as a central feature of RA pathology [12–14].

At the transcriptional level, reduced induction of *ATP5B*, *LDHA* and *DLD* may suggest a loss of metabolic flexibility rather than a defect in a single pathway. This pattern resembles metabolic inflexibility observed in heart failure with reduced ejection fraction (HFpEF), a condition more common in RA patients [53]. This is further supported by the lack of response to treatment. While healthy cardiomyocytes show increased metabolic activity with adalimumab, this effect is absent in snRA, consistent with reduced clinical response to anti-TNF therapy in seronegative patients [54]. Together with IL-6–mediated *KCNH2* downregulation, this defines a combined metabolic–electrical vulnerability in snRA, distinct in mechanism but converging functionally with the electrical phenotype of spRA.

### 4.5. Therapeutic implications

Neither anti-rheumatic agent restored normal cardiomyocyte function, but their effects differed. Adalimumab increased metabolic activity in healthy cells and partially in spRA, but had no effect in snRA. This suggests that TNF-α inhibition alone may not restore mitochondrial function in seronegative disease, which may contribute to persistent cardiovascular risk despite therapy [55]. Tofacitinib showed a different pattern. It increased *KCNJ3* expression in healthy cells but failed to induce this response in RA cardiomyocytes, where levels remained reduced. As *KCNJ3* channels stabilize membrane potential, this further weakens repolarization reserve in RA. Combined with impaired TNF-α signalling, this represents a cumulative loss of protective mechanisms. This may help explain the increased rate of major adverse cardiovascular events observed in the ORAL Surveillance trial [34]. These findings support ECG monitoring in patients receiving JAK inhibitors, consideration of serotype in treatment decisions and further *in vivo* testing of the proposed mechanism.

### 4.6. Limitations and future directions

Several limitations should be considered when interpreting these findings. hiPSC-derived cardiomyocytes retain an immature phenotype compared with adult cells and variability between lines persists despite current maturation strategies. The 24-hour exposure window captures early responses but does not reflect longer-term remodelling. In addition, ionic currents were not directly measured and the observed dissociation between single-cell and multicellular effects requires further mechanistic validation. The APD ratio applied here is not directly equivalent to classical triangulation indices, which may limit comparability. Finally, the molecular basis of impaired potassium channel inducibility remains unresolved and will require investigation at the chromatin level. Despite these limitations, the multi-modal approach - combining metabolic, electrophysiological and transcriptional analyses - provides a coherent view of RA-related cardiomyocyte dysfunction. Future clinical studies stratified by autoantibody status will be essential to test whether these mechanisms translate into distinct cardiovascular risk profiles.

## 5. Conclusions

This study defines a dual pattern of cardiomyocyte vulnerability in RA. Seropositive cells show primarily electrical dysfunction, while seronegative cells combine metabolic and electrical abnormalities. Despite these differences, both converge on reduced repolarisation reserve driven by impaired potassium channel regulation. This identifies the cytokine - potassium channel axis as a central mechanism linking inflammation to cardiovascular risk in RA. The observed limitations of current screening approaches and the impaired response to therapy highlight the need for serotype-aware strategies in both risk assessment and treatment. More broadly, patient-derived hiPSC cardiomyocytes provide a powerful platform for studying inflammation-driven cardiac dysfunction and may be extended to other chronic inflammatory diseases.

## CRediT authorship contribution statement

**Jan Wolnik:** Conceptualization, Data curation, Formal analysis, Investigation, Methodology, Software, Validation, Visualization, Writing – original draft, Writing – review & editing. **Patrycja Adamska:** Investigation, Methodology, Visualization, Writing – review and editing. **Aleksandra Oleksy:** Investigation, Methodology, Project administration, Software, Writing – review and editing. **Anna Magdalena Sanetra:** Data curation, Formal analysis, Investigation, Methodology. **Katarzyna Palus-Chramiec:** Data curation, Formal analysis, Investigation, Methodology. **Marian Henryk Lewandowski:** Resources, Supervision. **Józef Dulak:** Conceptualization, Resources, Supervision. **Monika Biniecka:** Conceptualization, Funding acquisition, Project administration, Resources, Methodology, Supervision, Writing – original draft, Writing – review & editing.

## Declaration of Generative AI and AI-assisted technologies in the writing process

AI-assisted technology was not used in the development of this manuscript, except for grammar and spelling checks.

## Funding

Study supported by the National Science Centre: OPUS 13 (UMO-2017/25/B/NZ5/02243 to MB and Sonata Bis 8 (UMO-2018/30/E/NZ5/00488 to MB) and by Horizon 2020 (841627-H2020-MSCA-IF to MB). The research with multi-electrode array (MEA) system used for electrophysiological analyses was supported by the National Science Centre grant SHENG-2 (2021/40/Q/NZ3/00165) to J.D.

## Declaration of competing interest

The authors disclose no conflict of interest.

## Acknowledgement

We would like to acknowledge Prof. Ursula Fearon and Prof. Douglas Veale for providing us FLS samples, as well as dr Marta Skoczyńska, Prof. Piotr Wiland and dr Grzegorz Kubiak for providing us PBMC samples. We also thank the late Ms. Agnieszka Andrychowicz-Róg and Ms. Joanna Uchto-Bajołek for their administrative assistance. We also express our sincere appreciation to the patients who participated in this study. Figures were created using Biorender.com.

## Data availability

The data that support the findings of this study and materials are available from the corresponding author upon reasonable request.

## Additional material provided

1. **Additional file 1: Supplementary Methods.** Detailed description of the experimental procedures.
2. **Additional file 2: Supplementary Table 1**. Donor demographic and clinical characteristics.
3. **Additional file 3: Supplementary Table 2**. Primary and secondary antibodies used for immunofluorescent staining.
4. **Additional file 4: Supplementary Table 3**. RT-qPCR primer sequences.
5. **Additional file 5: Supplementary Table 4**. Definitions and physiological interpretation of action potential parameters.
6. **Additional file 6: Supplementary Fig. 1. Validation of patient-derived hiPSC and CMs. (A)** Karyotype: G-banded metaphase analysis confirming normal karyotype with no structural or numerical aberrations. **(B) Pluripotency** markers: Immunofluorescence images of undifferentiated hiPSC colonies stained for NANOG (green), TRA-1-60 (green) and TRA-1-81 (green); nuclei counterstained with DAPI (blue). Scale bar: 100 µm. **(C)** Representative gating strategy for cardiac troponin T-positive (cTnT⁺) cardiomyocytes on day 21 of differentiation, prior to MACS purification. Sequential gating: P1 — cell population gate on FSC-A/SSC-A (debris exclusion); P2 — singlet discrimination on FSC-H/FSC-W (doublet exclusion); P3 — live cells (DAPI⁻); P4 — viable single cells progressed to cTnT analysis; P5 — cTnT⁺ cardiomyocytes identified on the Alexa Fluor 488. The representative line shown contained 76.8% cTnT⁺ cells prior to MACS enrichment, consistent with typical Wnt-modulation differentiation yields. Only post-MACS hiPSC-CM populations were used in downstream functional assays.
7. **Additional file 7: Supplementary Fig. 2. Additional patch-clamp parameters.** Maximum upstroke velocity (**top**), AP peak amplitude (**middle**), AP through (**bottom**) in HC (blue), spRA (red), snRA (green). Plots display the distribution of individual cell measurements, with median (solid line) and interquartile range (dashed lines). Kruskal–Wallis with Dunn’s: *p < 0.05; **p < 0.01; ***p < 0.001; ****p < 0.0001.
8. **Additional file 8: Supplementary Fig. 3. Representative MEA recordings of hiPSC-CMs exposed to pro-inflammatory cytokines.** Spontaneous electrophysiological activity of hiPSC-CMs was recorded on 24-well Multiwell-MEA plates before and 24 h after stimulation with pro-inflammatory cytokines IL-1β, IL-6 and TNF-α. For each condition, representative traces are shown at baseline (Day 0, background activity prior to stimulation) and after 24 h (Day 1, post-stimulation). Time-matched unstimulated wells (control) served as the reference for culture-time-related changes. **Top row:** averaged extracellular spike waveform centred on the depolarisation peak (±10 ms), reflecting the morphology of the fast depolarisation phase of the field potential. **Middle row:** full field potential (FP) waveform across the depolarisation–repolarisation cycle, used for QT/field potential duration (FPD) analysis (start offset 100 ms, stop offset 1 000 ms, noise suppression 10 ms, peak factor 150). Green markers indicate QT detection points obtained by automated detection and subsequently verified manually for every recording. **Bottom row:** continuous segment of the raw spontaneous beating signal from a representative electrode, illustrating beat rate, rhythm regularity and amplitude consistency across successive contractions over the 5-minute recording window. Comparison of Day 0 vs. Day 1 traces within each condition illustrates the impact of individual cytokines on cardiomyocyte electrophysiology, including changes in spike morphology, FPD and beating rhythm.

## Supplementary Methods

### Generation of hiPSC lines from PBMCs and FLS

Peripheral blood mononuclear cells (PBMCs) were obtained from whole blood using BD Vacutainer Mononuclear Cell Preparation Tubes (BD Biosciences). Isolated cells were maintained in suspension culture for 4 days in StemPro-34 SFM medium (Thermo Fisher Scientific) supplemented with 2 mM L-glutamine (Thermo Fisher Scientific), 100 ng/mL SCF (PeproTech), 100 ng/mL FLT-3 (PeproTech), 20 ng/mL IL-3 (PeproTech), and 20 ng/mL IL-6 (PeproTech). Synovial biopsies were enzymatically processed for FLS isolation using 1 mg/mL collagenase type I (Worthington Biochemical) in RPMI-1640 medium (Thermo Fisher Scientific) for 4 h at 37 °C. Following digestion, the resulting cell suspension was plated in RPMI-1640 medium supplemented with 10% FBS, 20 mM HEPES, 100 U/mL penicillin/streptomycin, and 0.25 µg/mL amphotericin B (all from Thermo Fisher Scientific). Reprogramming was performed by transducing PBMCs (200 000 cells) and FLS (50 000 cells) with Sendai viral vectors using the CytoTune-iPS 2.0 Sendai Reprogramming Kit (Thermo Fisher Scientific) in complete StemPro-34 SFM medium or complete FLS medium, respectively. After 24 h, the medium was replaced and cells were maintained under standard culture conditions. PBMC-derived cells were transferred onto Geltrex-coated wells 3 days post-transduction, whereas FLS-derived cells were plated after 7 days. From day 7 onward, cultures were maintained in E8 medium (Thermo Fisher Scientific) with daily medium exchange. Around day 15, individual colonies were manually picked and transferred to separate wells. Karyotype integrity of iPSCs was verified by G-banding (GTG-450, 15 mitoses per sample) by the Kariogen laboratory (Kraków, Poland).

### Characterisation of hiPSC pluripotency and trilineage differentiation potential

The pluripotent phenotype of each generated hiPSC line was verified through detection of pluripotency-associated markers using immunofluorescence, following the protocols detailed below. The capacity of the generated lines to differentiate into derivatives of all three germ layers was assessed using two complementary approaches: embryoid body (EB) formation followed by spontaneous differentiation *in vitro*, and teratoma formation *in vivo*.

### Embryoid body formation

hiPSCs were dissociated and plated at a density of 5 000 cells per well into U-bottom, non-adhesive 96-well plates in Essential 6 medium (E6; Thermo Fisher Scientific) containing polyvinyl alcohol (4 mg/mL; Sigma-Aldrich) and the ROCK inhibitor Y-27632 (10 µM; Abcam). After 24 h, the medium was exchanged for fresh E6 without supplements, and aggregates were cultured for a further 7 days to allow EB maturation. EBs were then transferred onto Geltrex-coated surfaces and left to spontaneously differentiate over a 14-day period.

### Teratoma formation assay

For the in vivo teratoma differentiation assay, 1 × 10^6^ hiPSCs per line were collected, rinsed in Ca^2+^/Mg^2+^-free PBS, and resuspended in a 3:2 (v/v) mixture of Matrigel (Merck) and PBS. The cell suspension was delivered by subcutaneous injection into the dorsal flank of female NOD SCID mice (strain NOD.Cg-*Prkdc*^scid^ *Il2rg*^tm1Wjl/SzJ^), which had been anaesthetised with inhaled isoflurane in a dedicated induction chamber. Animals were housed under specific pathogen-free conditions and received standard chow and water ad libitum. Body mass was recorded weekly and tumour dimensions were monitored on alternate days until reaching a volume of at least 1 cm^3^. Twelve weeks post-injection, mice were sacrificed by CO_2_ inhalation followed by cervical dislocation. Excised teratomas were fixed in formalin, paraffin-embedded, sectioned, and stained with haematoxylin and eosin for histopathological evaluation of tissues representative of the three germ layers. All animal experiments complied with Directive 2010/63/EU and were approved by the Second Local Ethical Committee for Animal Research in Kraków, Poland (approval no. 131/2021).

### Cardiac differentiation

Cardiac differentiation was induced through temporal modulation of the Wnt/β-catenin signalling pathway using sequential exposure to small-molecule agonists and antagonists. In the initial phase, mesodermal commitment was initiated via inhibition of GSK3β with CHIR-99021, thereby activating canonical Wnt signalling. Subsequent specification toward cardiac mesoderm was achieved through Wnt pathway inhibition using IWR-1. Undifferentiated hiPSCs were seeded at a density of 6 × 10^4^ cells per well in 24-well plates in E8 medium supplemented with 10 µM Y-27632 ROCK inhibitor for the first 24 h to enhance post-plating survival. Upon reaching ∼90% confluence (defined as day 0), the medium was replaced with RPMI-1640 supplemented with 2% B-27 minus insulin (RPMI/B27−ins; Thermo Fisher Scientific) containing 6 µM CHIR-99021 (Sigma-Aldrich). After 24 h (day 1), the medium was refreshed with CHIR-free RPMI/B27−ins. On day 3, cells were treated with IWR-1 (Sigma-Aldrich) at 4–7 µM for 48 h, with the optimal concentration determined empirically for each hiPSC line. Following this, cultures were maintained in compound-free RPMI/B27−ins until day 7. From day 7 onward, the medium was switched to RPMI-1640 supplemented with 2% B-27 (RPMI/B27; Thermo Fisher Scientific), with medium changes every 72 h until day 21. At this stage, differentiated monolayers were dissociated into single-cell suspensions using the Multi-Tissue Dissociation Kit 3 (Miltenyi Biotec) and subjected to magnetic-activated cell sorting (MACS). Cardiomyocytes were enriched to >90% cTnT^+^ purity using the PSC-Derived Cardiomyocyte Isolation Kit (Miltenyi Biotec). Purified cells were replated onto Geltrex-coated substrates appropriate for downstream applications. Functional assays were performed between days 30 and 45 post-differentiation.

### Mitochondrial function and oxidative stress assays

Unless otherwise specified, fluorescent probes used for oxidative stress and mitochondrial analyses were prepared in RPMI-1640 medium supplemented with B-27 (with or without phenol red, as indicated) and applied to live cells for 30 min at 37 °C. The probes and working concentrations were as follows: CellROX Deep Red (5 µM; in phenol-red-free RPMI-1640 + B-27), MitoSOX Red (5 µM; in RPMI-1640 + B-27), and MitoTracker Green FM (100 nM; in RPMI-1640 + B-27). After staining, cells were rinsed with PBS and dissociated with the Multi-Tissue Dissociation Kit 3 (Miltenyi Biotec). Flow cytometry data were acquired on an LSR Fortessa instrument (BD Biosciences) using FACSDiva software and further analyzed in FlowJo.

### Cytoplasmic and mitochondrial ROS detection

Oxidative stress was quantified in hiPSC-CMs seeded at 100 000 cells per well on Geltrex-coated 24-well plates. After a 48 h stabilisation phase followed by the designated stimulation, cells were labelled with either CellROX Deep Red or MitoSOX Red (both Thermo Fisher Scientific) to assess total intracellular or mitochondrial superoxide levels, respectively. CellROX Deep Red undergoes oxidation by reactive oxygen species and binds nucleic acids, thereby providing a readout of global cellular oxidative burden, whereas MitoSOX Red is selectively targeted to mitochondria and reports superoxide generated within the matrix. Tert-butyl hydroperoxide (TBHP) was included in each experiment as a positive control to confirm probe responsiveness.

### Mitochondrial mass

Mitochondrial content was estimated in parallel cultures seeded and stimulated under identical conditions. After stimulation, cells were labelled with MitoTracker Green FM (Thermo Fisher Scientific), a cell-permeant dye that accumulates within mitochondria in a membrane-potential-independent manner and covalently attaches to matrix proteins. The resulting fluorescence intensity served as a surrogate indicator of mitochondrial content per cell.

### Mitochondrial membrane potential (ΔΨm)

Changes in ΔΨm were evaluated using two complementary dyes: tetramethylrhodamine methyl ester (TMRM) and JC-1 (both Thermo Fisher Scientific). For TMRM-based analysis, hiPSC-CMs were seeded at 100 000 cells per well on 24-well plates and subjected to the appropriate 24 h stimulation. Cells were then dissociated, resuspended in culture medium containing 100 nM TMRM, and incubated for 30 min protected from light before acquisition on the LSR Fortessa. For the JC-1 assay, 10 000 cells per well were plated in black-walled 96-well plates (PerkinElmer) and cultured for 72 h. Cells were stained with JC-1 (10 µg/mL, 10 min, in the dark), washed, and fluorescence was recorded on an M200 plate reader (TECAN) at excitation/emission wavelengths of 488/530 nm (monomers) and 488/590 nm (J-aggregates); results are reported as the ratio of red (aggregate) to green (monomer) fluorescence. The protonophore FCCP was used as a positive control for mitochondrial depolarisation.

### Holotomography imaging

Subcellular distribution of the fluorescent probes was examined by live-cell holotomography on an HT-X1 microscope (Tomocube) at 40× magnification, combining quantitative phase imaging (QPI) with fluorescence microscopy. The QPI component captures refractive-index maps of intracellular structures without the need for fixation or additional labelling, while the fluorescence channels allow simultaneous visualisation of probe signals, preserving cell viability during extended acquisition. hiPSC-CMs were plated at low density (50 000 cells per glass-bottomed holotomography dish) and cultured for 48 h. Immediately before imaging, cells were incubated with a mixture of the probes described above for 30 min at 37 °C, washed with ion-free PBS, and acquired simultaneously in holotomography and fluorescence modes to enable spatial co-localisation analysis of mitochondrial and ROS signals.

### Flow cytometry characterization of cardiomyocytes

hiPSC-CMs were dissociated using EDTA (Invitrogen) or the Multi-Tissue Dissociation Kit 3 (Miltenyi Biotec). Single-cell suspensions were pelleted, fixed in 4% paraformaldehyde (Santa Cruz Biotechnology) for 20 min at room temperature, and rinsed twice in PBS containing 2% FBS. Membrane permeabilisation was performed with 0.1% Triton X-100 (BioShop Canada Inc.) for 30 min at room temperature. CMs were incubated with primary antibody against cardiac troponin T for 1 h at room temperature and with fluorophore-conjugated secondary antibodies for 30 min at room temperature in the dark (Supplementary Table 2). Nuclei were counterstained with DAPI (0.2 µg/mL; Sigma-Aldrich) prior to acquisition. Data were collected on an LSR Fortessa cytometer (BD Biosciences) using FACSDiva software and analysed in FlowJo. Gating was performed on singlet, DAPI-positive events and compared to unstained controls.

### Immunofluorescence staining

The presence of specific markers was evaluated by immunofluorescence microscopy in hiPSCs, embryoid bodies (EBs), and MACS-purified hiPSC-CMs. Cells were plated onto 10 mm glass coverslips pre-coated with Geltrex within 24-well plates. After the experimental endpoint, samples were fixed in 4% paraformaldehyde for 20 min at room temperature, rinsed three times in Ca^2+^/Mg^2+^-free PBS, and permeabilised with 0.1% Triton X-100 for 10 min. All samples were subsequently blocked in PBS containing 3% bovine serum albumin (BSA; BioShop Canada) for 1 h at room temperature. Primary antibodies diluted in blocking buffer were applied overnight at 4 °C (Supplementary Table 2). The following day, coverslips were washed three times in PBS, incubated with fluorophore-conjugated secondary antibodies for 1 h at room temperature in the dark (Supplementary Table 2), and counterstained with DAPI (0.2 µg/mL; Sigma-Aldrich). Mounted coverslips (Dako Fluorescent Mounting Medium, Agilent) were imaged on an Axio Observer inverted fluorescence microscope (Carl Zeiss, Oberkochen, Germany).

### RT-qPCR

RNA yield and spectral purity (A260/280 and A260/230 ratios) were verified on a NanoDrop 1000 spectrophotometer (Thermo Fisher Scientific). Each qPCR reaction (final volume 15 µL) contained 7.5 µL of AceQ Universal SYBR qPCR Master Mix (Vazyme), 0.4 µL each of forward and reverse primer (final working concentration 0.25 µM per primer), 4.7 µL of nuclease-free water, and 2 µL of cDNA template. Amplification was carried out on a StepOnePlus real-time thermocycler (Applied Biosystems) running StepOne v2.3 software under the default cycling programme: initial denaturation at 95 °C for 5 min, followed by 40 cycles of 95 °C for 10 s, 60 °C for 30 s, and 72 °C for 30 s, with a final melt curve analysis. Each sample was analysed in technical triplicate. Primer sequences are provided in Supplementary Table 3.

### Patch-clamp specifications

#### Recording solutions

Both intracellular and extracellular solutions were filtered through 0.22 µm sterile filters before use. The extracellular solution was freshly prepared every two days, whereas the intracellular solution was prepared in bulk, stored as single-use aliquots at −20 °C, and a fresh aliquot was thawed each day prior to recording. Solution osmolarity was verified using a Vapro 5520 osmometer (Wescor) and adjusted with sucrose if required. Solutions were equilibrated to 37 °C in a water bath prior to recording.

#### Cell selection and quality control

Only single, isolated cardiomyocytes displaying stable membrane potential, clear responses to depolarising stimuli, and ventricular-like action potential morphology were included in the analysis. Inclusion criteria comprised: (i) plateau-containing phase 2; (ii) maximum diastolic potential more negative than −60 mV; (iii) AP amplitude greater than 70 mV; (iv) regular firing pattern with coefficient of variation of inter-beat interval below 30%; and (v) stable series resistance throughout the recording. Cells displaying spontaneous pacemaker-like phase 4 depolarisation, irregular firing patterns, or signs of mechanical damage or membrane rupture were excluded prior to data analysis.

#### Action potential parameter definitions

Six action potential parameters were extracted per cell using Signal software (CED): (i) maximum upstroke velocity (dV/dt max), the maximum first-derivative value of the voltage trace during phase 0 depolarisation, expressed in mV/ms; (ii) AP peak, the most positive membrane potential attained during the action potential, expressed in mV; (iii) AP through, the most negative membrane potential reached between consecutive action potentials during phase 4, expressed in mV; (iv) APD20, the time interval from AP peak to 20% repolarisation, expressed in ms; (v) APD50, the time interval from AP peak to 50% repolarisation, expressed in ms; (vi) APD90, the time interval from AP peak to 90% repolarisation, expressed in ms. The APD ratio (30–40 / 70–80) was additionally calculated as (APD40 − APD30) / (APD80 − APD70) to provide a shape-sensitive index. Values greater than 1 indicate proportionally extended early-phase repolarisation consistent with action potential triangulation, whereas values less than 1 indicate plateau-dominant ventricular-like morphology.

### MEA recording and analysis settings

Unstimulated hiPSC-CMs were seeded onto Geltrex-coated 24-well MEA plates (Multi Channel Systems), ensuring that only the electrode area was populated by cells, in 0.5 mL of RPMI-1640 medium supplemented with B-27. After 24 h, an additional 0.5 mL of fresh medium was added, and the cells were cultured for a further 3 days to allow the establishment of cell-cell connections, monolayer formation, and synchronisation of contractile activity. Spontaneous electrophysiological activity was recorded using a Multiwell-MEA system (Multi Channel Systems) at 37 °C and analysed using Multiwell-Screen software. Beating wells were selected for the experiments with a minimum of 3 wells per condition. Prior to stimulation, background electrophysiological activity was recorded for 5 min at 37 °C with a sampling rate of 20 000 Hz, high-pass filter 1 Hz, and low-pass filter 3 500 Hz. Subsequently, cells were stimulated by removing 0.2 mL of medium and adding 5× concentrated stimulants. After 24 h of stimulation, the electrophysiological signal was measured again with identical settings. Data were collected and analyzed for all active electrodes. Raw signals were processed using the Multiwell-MEA-Analyser software. A full 5-minute recording with activated Notch filter was used for analysis of each well in which at least one electrode measured constant, unwavered contractility throughout the recording time. QT analysis was performed with a start offset of 100 ms and a stop offset of 1 000 ms, noise suppression of 10 ms, and peak factor 150; other settings were left at default values. Each QT detection point was manually verified after automatic detection to ensure reliable results. Results were normalized to the means of background electrophysiological activity for each donor and compared to unstimulated control wells.

**Supplementary Table 1.** Donor demographic and clinical characteristics.

| No | Line ID | Source | Diagnosis | RF/ACPA status | Age | Sex | DAS-28 |
| --- | --- | --- | --- | --- | --- | --- | --- |
| 1 | 3 | PBMC | HC | - | 35 | F | - |
| 2 | 4 | PBMC | HC | - | 45 | M | - |
| 3 | 5 | PBMC | HC | - | 45 | F | - |
| 4 | 13 | PBMC | RA | sero positive | 64 | M | 5.52 |
| 5 | 14 | PBMC | RA | sero positive | 52 | F | 5.60 |
| 6 | 1 | FLS | RA | sero positive | 55 | M | 6.18 |
| 7 | 2 | FLS | RA | sero positive | 58 | F | 5.83 |
| 8 | 15 | PBMC | RA | sero negative | 61 | F | 7.16 |
| 9 | 17 | PBMC | RA | sero negative | 61 | F | 5.12 |
| 10 | 6 | FLS | RA | sero negative | 62 | F | 5.2 |
| 11 | 7 | FLS | RA | sero negative | 59 | M | 5.62 |

**Supplementary Table 2.**
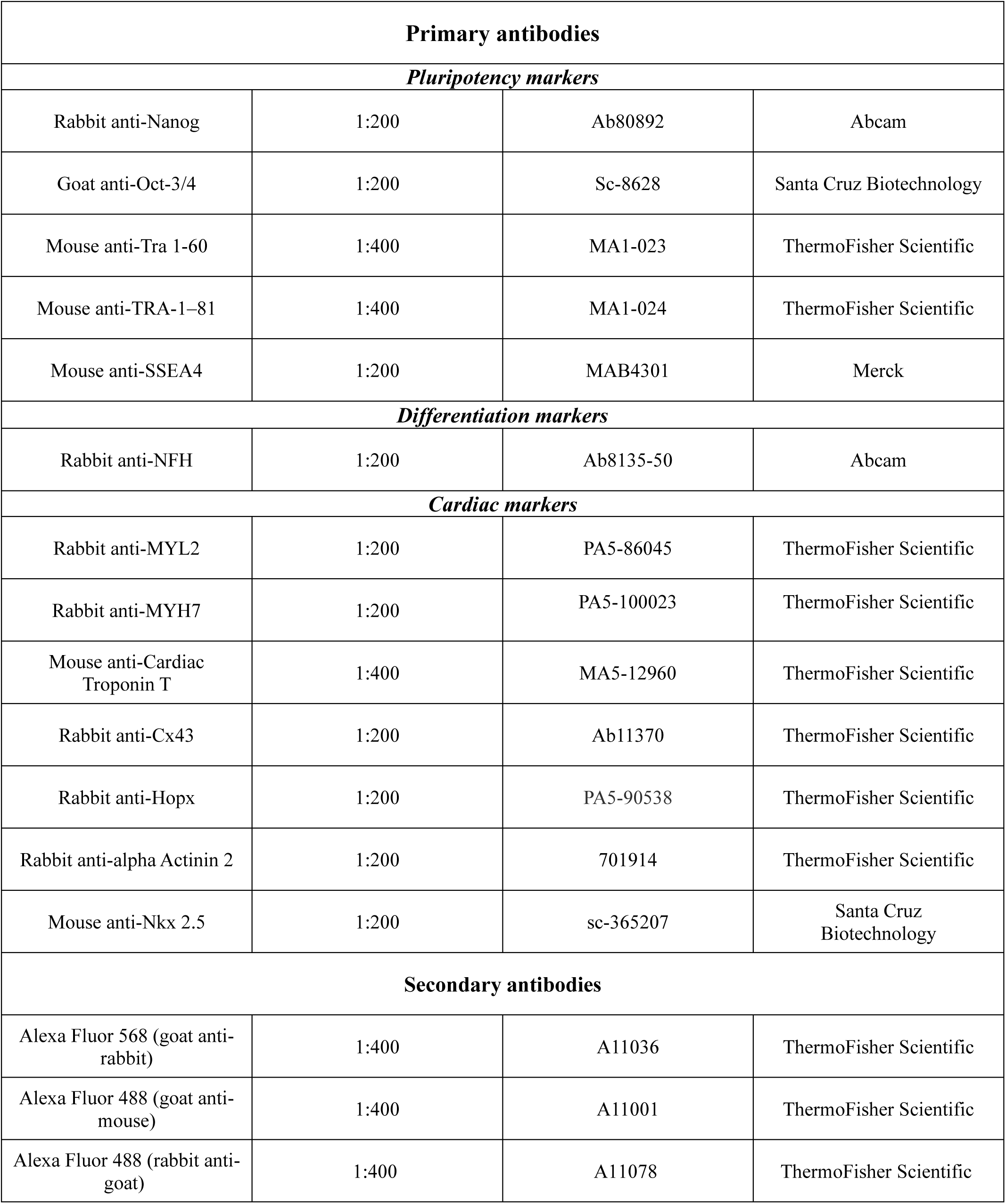
Primary and secondary antibodies used for immunofluorescent staining.

**Supplementary Table 3.**
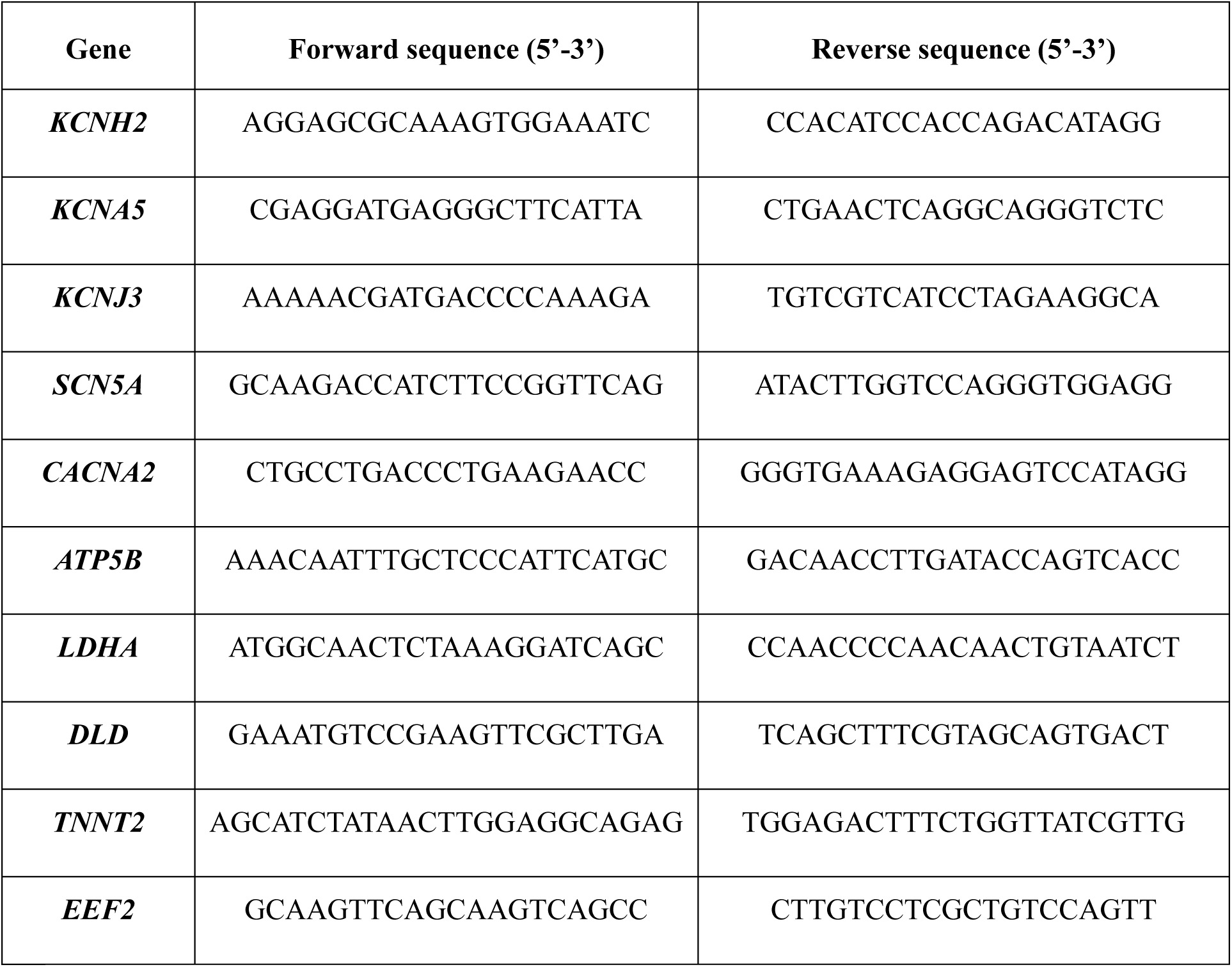
RT-qPCR primer sequences.

**Supplementary Table 4.** Definitions and physiological interpretation of action potential parameters.

| Parameter | Definition | Physiological interpretation |
| --- | --- | --- |
| ***Maximum upstroke velocity ($dV/dt_{max}$)*** | Maximum rate of membrane potential change during phase 0 depolarisation, calculated as the peak first derivative of the voltage trace during the rising phase | Surrogate of sodium channel (Nav1.5, *SCN5A*) function; higher values indicate stronger Na<sup>+</sup> influx during rapid depolarisation |
| ***Action potential peak (AP peak)*** | Most positive membrane potential reached during the action potential | Reflects the balance between Na<sup>+</sup> influx and opposing early repolarisation currents |
| ***AP trough*** | Most negative membrane potential between consecutive action potentials (phase 4) | Determined primarily by IK1 (*KCNJ2/KCNJ12*), IK<sub>ACh</sub> (*KCNJ3*), and Na<sup>+</sup>/K<sup>+</sup>-ATPase activity; more negative values indicate greater electrical stability |
| ***APD<sub>20</sub>*** | Time from AP peak to 20% repolarisation | Reflects early repolarisation currents, mainly Ito (Kv4.3) and IK<sub>ur</sub> (*KCNA5*) |
| ***APD<sub>90</sub>*** | Time from AP peak to 90% repolarisation | Reflects near-complete repolarisation; cellular correlate of QT interval; governed by IK<sub>r</sub> (*KCNH2*), IK<sub>s</sub> (*KCNQ1*), and ICa,L (*CACNA1C*) |
| ***APD ratio (30–40 / 70–80)*** | Ratio of repolarisation intervals between 30–40% and 70–80% repolarisation | Index of action potential shape; values >1 indicate relatively enhanced early repolarisation, while values <1 indicate a plateau-dominant morphology |

**Supplementary Fig. 1.**
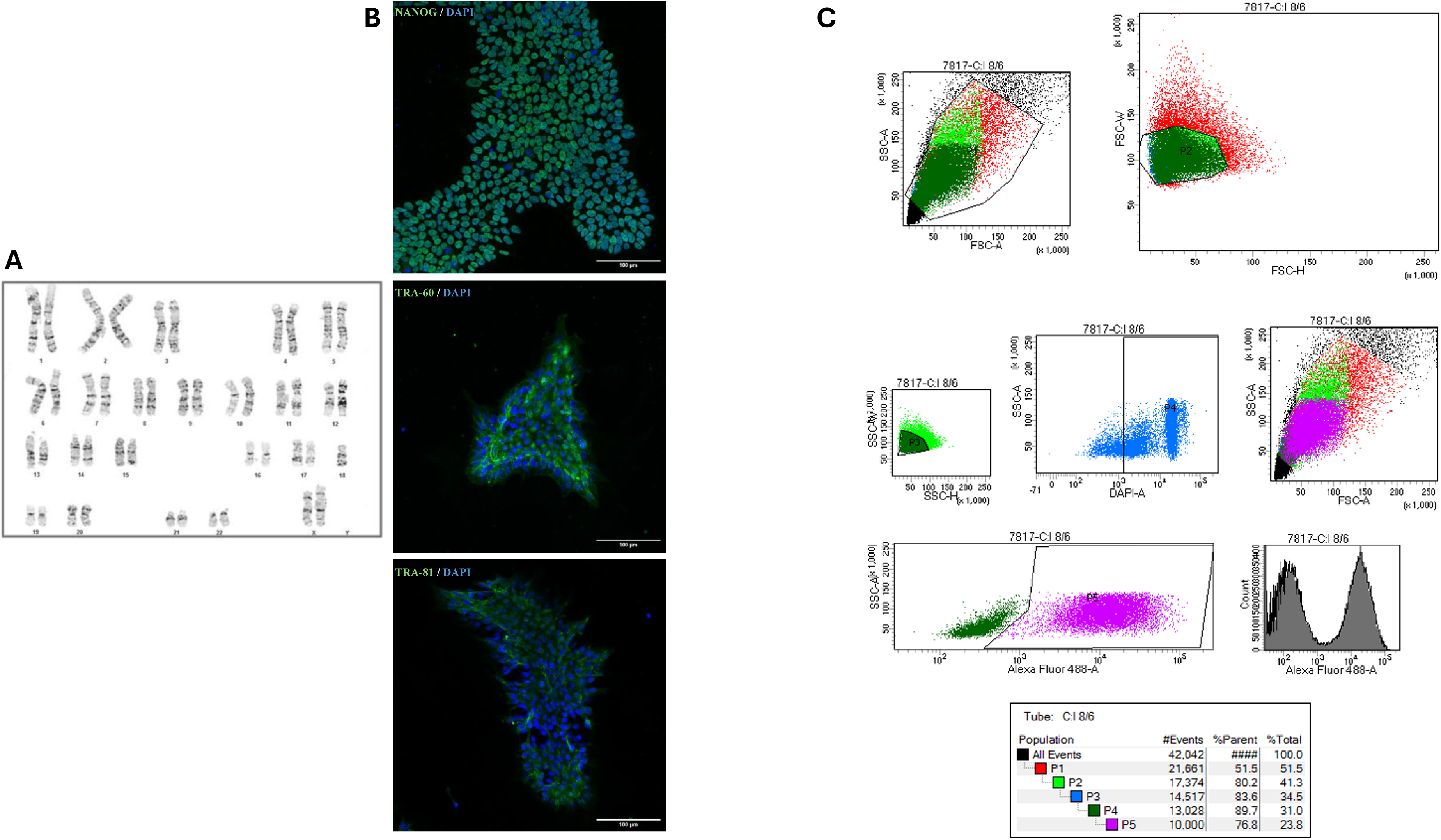

**Supplementary Fig. 2.**
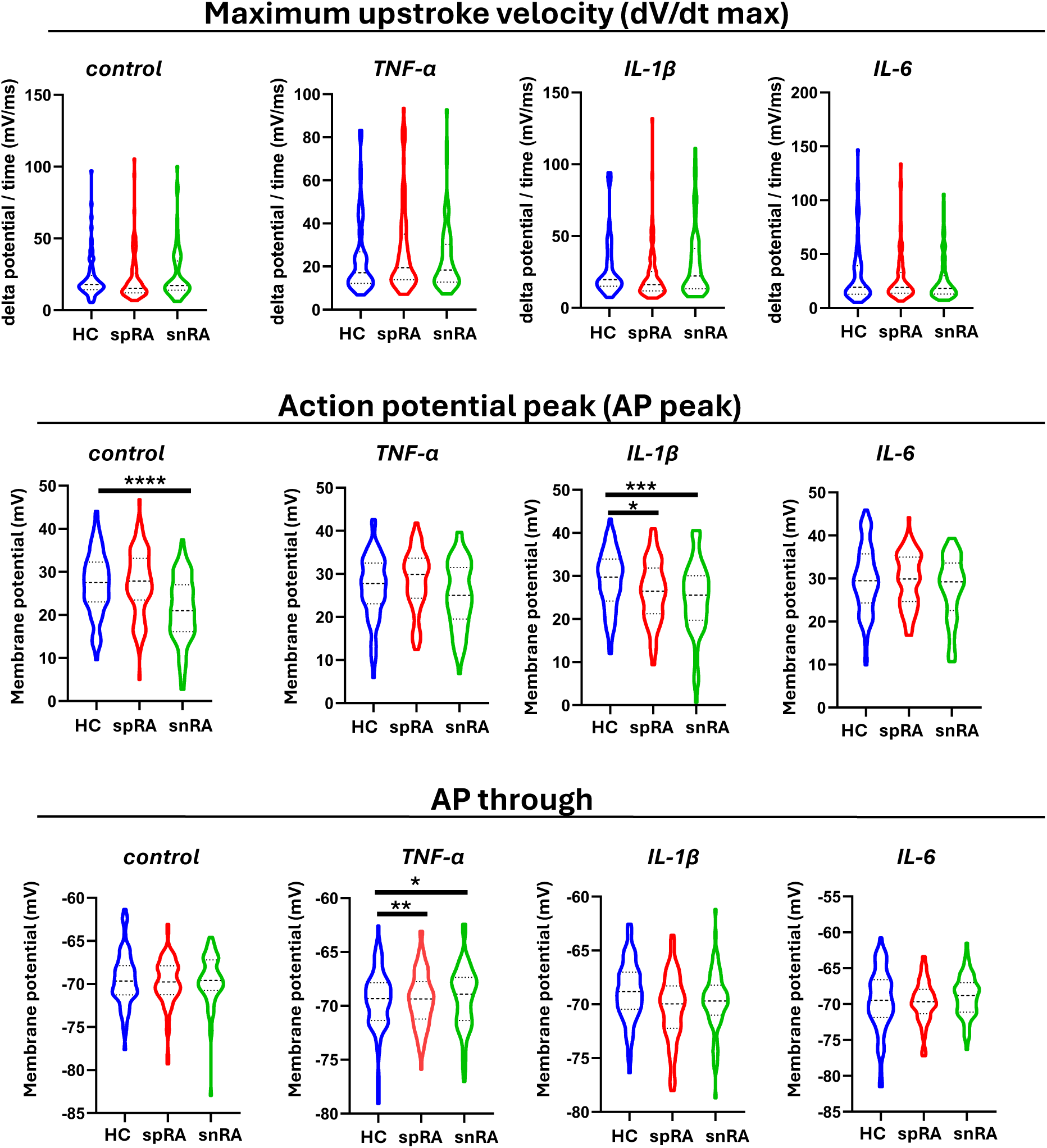

**Supplementary Fig. 3.**
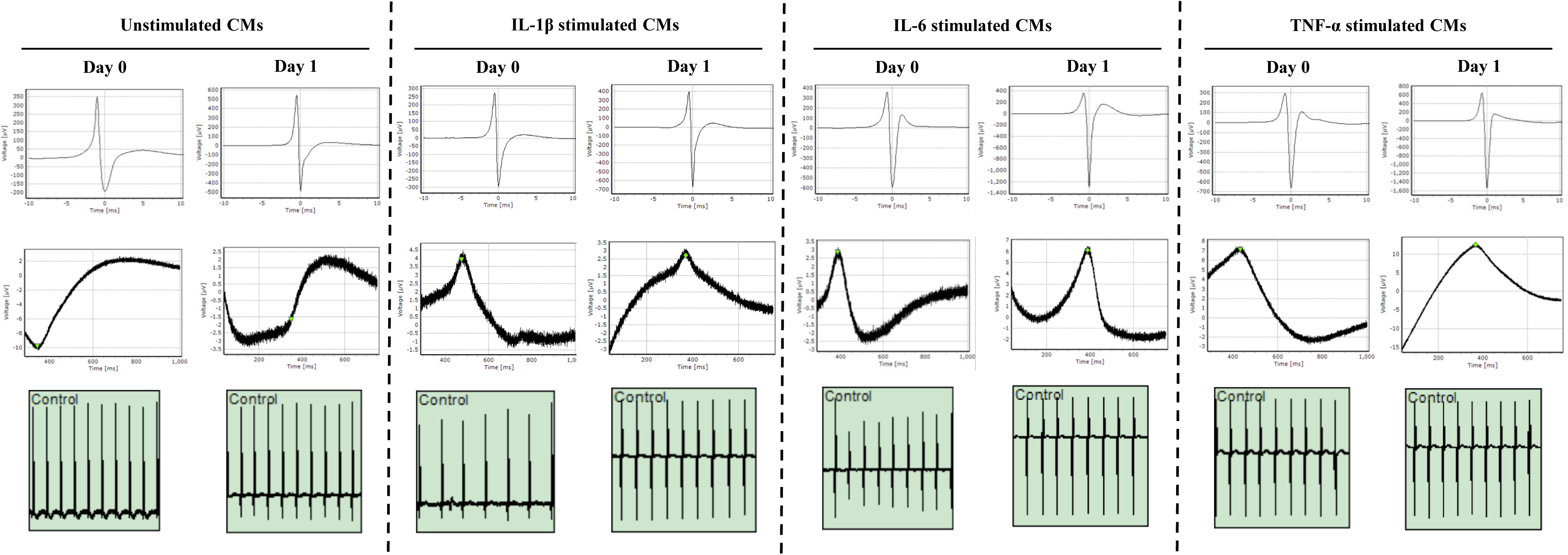

## Notes

### Competing Interest Statement

The authors have declared no competing interest.

